# Medial temporal lobe encodes cognitive maps of real-world social networks

**DOI:** 10.1101/2025.03.18.643925

**Authors:** Yi Yang Teoh, Jae-Young Son, Alice Xia, Apoorva Bhandari, Oriel FeldmanHall

## Abstract

People routinely navigate their complex social networks^1^: From gossiping strategically with others^2–4^ to brokering connections between siloed groups^5,6^, our ability to make adaptive social choices hinges on whether we can construct useful mental representations of the social ties within our communities^7^. While decades of neuroscience research have shown that the medial temporal lobe encodes cognitive maps of physical^8–10^ or conceptual space^11^, how the brain represents our social networks in the wild to solve social problems remains unknown. By combining computational models with functional neuroimaging and longitudinal measurement of an evolving and densely interconnected real-world human network (N=187), we show that the entorhinal cortex encodes a cognitive map of the long-range connectivity between pairs of network members. This social map reflects the particular demands of social navigation and is specifically formatted to encode the simultaneous connectivity between network members, which critically enables tracking how information diffuses across the network. Moreover, the strength of its encoding in the entorhinal cortex aids in brokering connections that improve cohesion within people’s social communities. Our results illuminate how a domain-general neural mechanism^12,13^ is tailored to prioritize the natural dynamics of social phenomena in order to support adaptive navigation through these highly complex environments.

## Main

To access the rich resources embedded in social networks and flourish^1^, people must learn to solve a variety of problems related to navigating complex social environments. From organizing a dinner party where no one feels left out to spreading pertinent gossip about an abusive co-worker, our ability to make adaptive choices hinges on whether we can construct a useful mental representation of the social relations within our communities. Different social problems demand different kinds of information—e.g., deep knowledge of social milieu to decide seating arrangements for an intimate dinner, versus inferences about social connections within the broader network that ensure gossip stays contained. Given the sheer number and variety of challenges people face in their social worlds, neural representations of social networks must be structured to enable maximal behavioral flexibility. How exactly does the brain solve this representational problem?

To meaningfully answer this question, we must examine how the brain naturally accomplishes this feat in large and dynamic real-world social networks. Although artificial social networks permit researchers exceptional experimental control during learning^14^, it is unknown whether representations of small artificial networks that lack many naturalistic qualities mirror how people spontaneously represent their own social networks in the wild. However, probing naturalistic representations of real-world networks introduces unique challenges: researchers cannot control how an individual observes their peers’ social interactions, and thus have no direct access to what people know about their social networks.

Advancements in computational modelling provide an elegant approach to addressing the challenges of studying natural social networks. Equipped with a minimal set of assumptions about the observations people can realistically make given the constraints of the real world, modelling allows us to synergistically draw on insights from relevant research on cognitive maps in controlled environments^15–20^ and apply them to far noisier real-world analogs.

One such insight is the critical role of the hippocampus (HC) and entorhinal cortex (EC) in constructing and maintaining map-like representations of relations between objects, locations, and concepts^8,10,21–26^. These same regions have also been implicated in the mapping of two-dimensional social hierarchies where people have to keep track of others’ power and competence^27,28^, further supporting the idea that the HC and EC play a fundamental role in representing relational information across domains. We might therefore expect similar neural computations in the HC and EC to support representations of our own large and complex social networks.

What relational information might these regions encode about social networks? One intuitive possibility is that people try to memorize a complete and veridical map of all the relationships they encounter in their social network. However, when we consider the size of real-world social networks, the immense cognitive demands of constructing and deploying such representations would render them highly inefficient^29,30^. Not only does brute-force memorization of all observed relations present a formidable cognitive challenge^31,32^, but information about others’ social relations is also not always apparent or directly observable^33^. Instead, recent work finds that people construct ‘fuzzy’ mental representations of others’ social relations^31,32,34,35^. These abstract representations of social networks emerge from a process of ‘chaining’ knowledge about pairwise social relations to infer the existence of indirect, multi-step connections between people. For example, if we know that Jane and Chris are friends, and that Chris and Zach are friends, then we might infer that Jane and Zach are also likely to be friends by abstracting over the known pairwise friendships and identifying the indirect connection between them^36,37^. By leveraging this simple mechanism of *multi-step abstraction*, people adeptly navigate a wide range of social situations—from generating inferences about unobserved but likely relationships within a network^31,32^, to tracking how gossip might spread across dense ties within a community^35^.

Given these considerations, it is unlikely that people construct a complete and veridical map of all social relationships they encounter. We instead hypothesize that people’s neural representations of their social networks likely reflect abstractions over whatever relations they have observed. Such abstraction would be consistent with research in the non-social domain in which the anterior HC and the EC appear to encode abstract representations of task environments, with the HC maintaining representations of multi-step connections between states in reinforcement learning tasks^18,19^, and the EC abstracting over these representations to encode more latent structural properties of the state-spaces^12,15,17,20^. However, whether the HC and/or EC encodes abstract representations of relational ties in the real world to support adaptive social navigation, remains wholly unknown.

It is also unclear what specific format these representations should take. Competing representational formats prioritizing different theoretical considerations have been proposed. On one hand, because each person maintains multiple relationships with many others, we might expect that an abstract representation of a social network tracks the absolute multi-step connectivity of each person to everyone else *independent* of their other connections within the network. Such a representational format—captured by Katz communicability (Katz)^35,38–41^—would be necessary if one wanted to anticipate the natural dynamics of information flowing through the network^35^. Gossip, for example, is often simultaneously transmitted across many social ties at once^2–4,42^, which demands a representation that accommodates variations in the overall connectedness of each person. On the other hand, representations of relationships in the network can also be structured in such a way that they quantify each person’s *relative* connectivity to all others in the network^31,32^. This representational format—known as the Successor Representation (SR)^43^—tracks the relative likelihood of people sharing multi-step connections with others in the network, which resembles the representations often used for navigating other types of environments^18,30,43,44^. Research illustrates that the SR captures how people sequentially navigate through these environments like grid worlds or physical space, tracking the relative probability that moving in one direction (over another) will lead you closer to an ultimate goal^20^. Such a representational format would limit accurate tracking of social transmission to cases where it unfolds sequentially, such as a note or physical object passed from one person to another over time. Although this does not align with the properties of social transmission (e.g., gossip), should we find evidence of an SR-like representation of real-world social networks, it could reflect a domain-general constraint on how highly complex relational information is encoded.

To close these fundamental gaps in our understanding of whether and how social networks are naturally—and spontaneously—represented in the HC and EC, we leveraged the unique context of first-year undergraduates leaving home and moving to college. This allowed us to examine the socio-cognitive maps people construct when embedded in a newly emerging and rapidly evolving social network. We tracked the emergence of the network (N=187) longitudinally, starting when students first arrived on campus, to the stabilization of friendships by the end of the academic year. Halfway through the year, once students established some initial friendships, we scanned a subset of the network (N=43) to interrogate whether HC and EC encode representations of one’s real social network. By developing and comparing a suite of computational models, we characterize the precise format in which the brain represents social relations in this naturalistic, ecologically valid, and personally relevant social network. Finally, we test whether these neurocognitive maps predict the ability to solve various social navigation problems, from asking whether these maps can track how information flows across the entire network to probing whether they enable the brokerage of friendships to foster greater social cohesion within people’s immediate social communities.

### People build imprecise cognitive maps of their real-world social network

We recruited 196 first-year undergraduates from three dormitories at Brown University and measured the ‘ground-truth’ structure of the friendship network by asking each participant to identify their friends amongst the other 195 participants multiple times throughout the year (Fig 1A; see Methods). After a second measurement of the network approximately one month into the academic year when students began to establish some initial friendships (N = 187), we recruited a subset of these network members (N=100) to complete a laboratory task probing participants’ mental representation of their social network. In this Network Knowledge Task, participants were presented with the name and photograph of a network member (target) alongside a list of names and photographs of all other network members (probes) and asked to judge whether each pair were friends.

**Fig 1.**
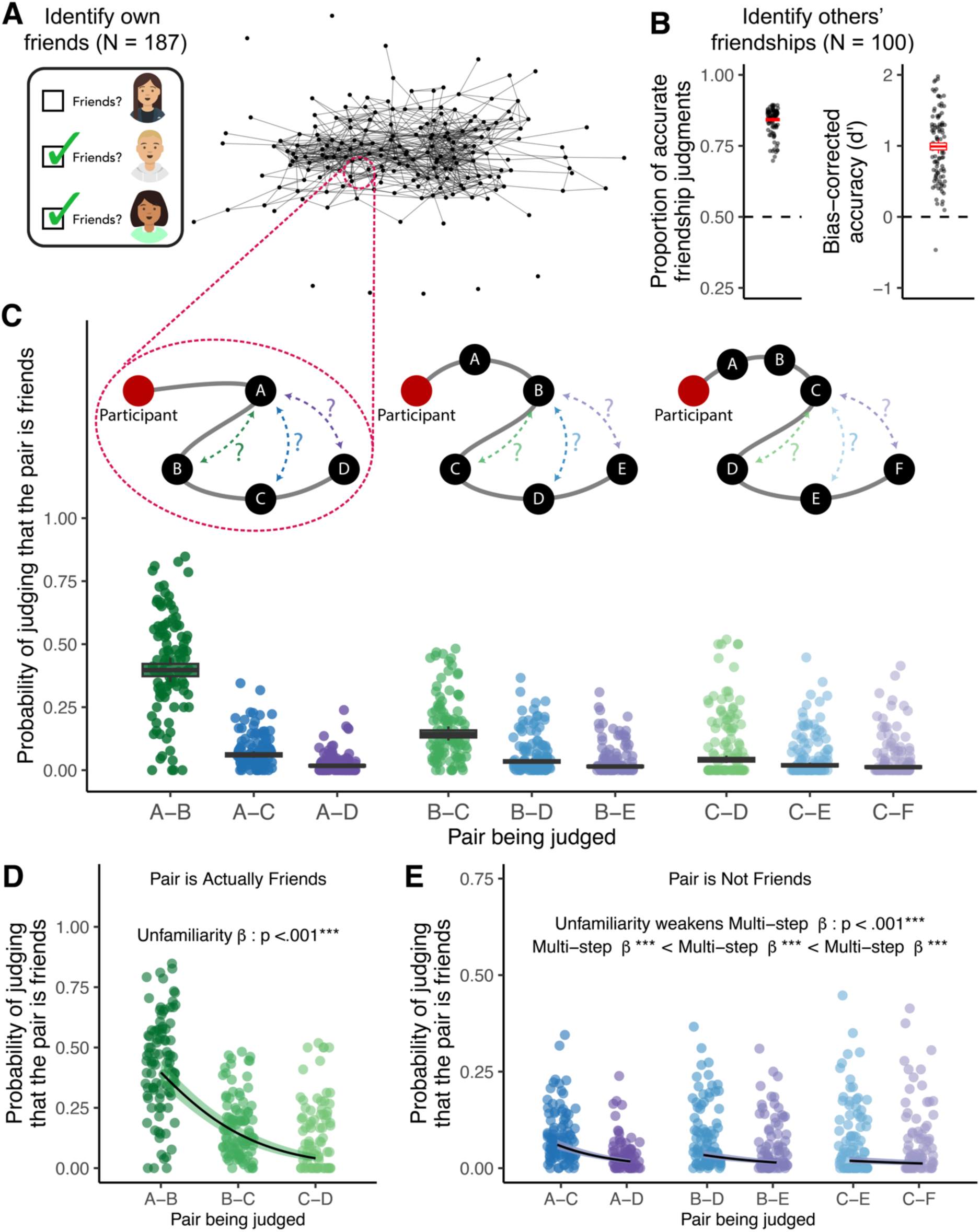
Friendship knowledge. **A) The friendship network constructed from mutually identified friendships.** In the graph (right), each node represents a network member who reported their own friendships in the friendship survey (left). Each edge between two nodes indicates a mutual friendship. Icons: getavataaars.com. **B) Accuracy of friendship judgments.** The two panels indicate (left) the proportion of accurate choices people made and (right) the bias-corrected accuracy, d-prime, which accounts for the relatively low base-rate of friendships within the network out of all possible dyadic relations that could exist. Each point represents a single participant’s data. The central line, upper and lower bounds, and whiskers of the boxplot indicate the estimated mean, within-participant std error, and 95% confidence interval from the general linear model. Dashed lines correspond to chance for each metric. **C) Friendship judgments reflect both direct and inferred knowledge.** Each dot represents the proportion of trials on which a participant indicated that a pair of network members were friends in the Network Knowledge Task conditional on whether the pair was actually friends (green), separated by a mutual friend (blue: friends-of-friends), or separated by three steps (purple: friends-of-friends-of-friends). These choices were further faceted out by participants’ familiarity with the pair as indicated by the visual schematics above the data plots: whether the participant was (left) personally friends with one of the members of the pair, (middle) separated by a mutual friend, or (right) separated by three steps from the pair (i.e., one of the pair’s members was a friend-of-a-friend-of-a-friend for the participant). Not all data is plotted here as there were very few trials in which the degree of separation—either within the pair or between the participant and the pair—exceeded three (6.45%). The central, line, upper and lower bounds, and whiskers of the boxplot indicate the estimated mean, within-participant std error and 95% confidence interval of response from a logistic mixed-effects regression fitted to all data respectively. **D) Friendship knowledge depends on familiarity with the pair.** This panel replots participants’ choices in the Network Knowledge Task in (c), but only for pairs of targets and probes that were actually friends, highlighting that people are less able to identify friends with whom they are less familiar. Each point represents the proportion of trials a participant indicated that the pair was friends. The black line and shaded region indicate the estimated mean and within-participant 95% confidence interval of responses from the fitted logistic mixed-effects regression. **E) Erroneous multi-step inferences.** This panel reproduces participants choices in the Network Knowledge Task in (c), but only for pairs of targets and probes that were not actually friends, highlighting that people are more likely to (incorrectly) infer friendship between dyads who are separated by two (blue) vs. three steps (purple). However, these multi-step inferences are modulated by the participant’s familiarity with the pair, such that greater unfamiliarity decreases the likelihood of making incorrect multistep inferences (unfamiliarity x multi-step interaction p < .001). Each point represents the proportion of trials a participant indicated that the pair was friends. The black line and shaded region indicate the estimated mean and within-participant 95% confidence interval of responses from the fitted logistic mixed-effects regression.***p< .001

We find that participants were able to identify whether other network members were friends with each other at above-chance accuracy (Chance = 0.50; Proportion correct responses: M = 0.843, 95% CI = [0.834, 0.850], z = 53.719, p < .001, Fig 1B). This was true even when using indices of accuracy that account for the low-base rate of friendships (i.e., only 14.5% of the dyads presented to participants were actually friends on average): d-prime confirms that people possess actual knowledge of friendships and were not simply responding randomly according to base rates (d-prime corresponding to chance responding = 0; M = 0.991, 95% CI = [0.894, 1.089], t(99) = 20.203, p < .001, Fig 1B).

However, as expected, participants also made systematic errors that illustrate (1) limitations in their knowledge about the network, and (2) the use of multi-step abstraction to make inferences about the existence of others’ friendships (Supplementary Table S1). First, even in the best-case scenario where a participant is friends with one of the people in the dyad (Fig 1C, pair A-B in dark green), the participant only accurately identifies ∼40% (M = 0.397, 95% CI = [0.349, 0.445]) of true friendships on average—a far cry from having complete knowledge of the network’s structure. This accuracy further degrades as a function of the participant’s familiarity with the pair being evaluated (unfamiliarity β = −1.370, 95% CI = [−1.562, −1.178], z = −14.012, p < .001; pair A-B vs pair B-C vs pair C-D, Fig 1D). If the participant is not directly friends with anyone in the dyad, but is instead separated by a mutual friend, the participant becomes less accurate (i.e., B-C: M = 0.143, 95% CI = [0.115, 0.171]), which decreases even further when separated by three-degrees (C-D: M = 0.041, 95% CI = [0.026, 0.055]).

Second, participants also erroneously endorse non-existent friendships in systematic ways that suggest they are doing some form of multi-step abstraction. When a participant evaluates whether their friend is friends with someone else—but the two are not in fact friends—the participant is more likely to incorrectly infer a friendship if the pair share mutual friends (triadic closure^37^: A-C: M = 0.061, 95% CI = [0.048, 0.073]) compared to if the pair are separated by three degrees (pair A-D: M = 0.030, 95% CI = [0.023, 0.038]; A-C vs A-D: Multi-step β = −1.295, 95% CI = [−1.554, −1.036], z = −9.810, p < .001, Fig 1E). These erroneous friendship endorsements reveal that participants use multi-step connectivity in the network to infer who is friends with whom.

This multi-step abstraction persists even if the participant is herself not directly friends with one person in the pair but is separated by mutual friends (B-D vs B-E: multi-step β = −0.878, 95% CI = [−1.048, −0.708], z = −10.110, p < .001) or three-degrees of friendship (C-E vs C-F: Multi-step β = −0.461, 95% CI = [−0.683, −0.239], z = −4.068, p < .001)—despite the overall frequency of these inferences decreasing as a function of being unfamiliar with the pair (difference in Multi-step β with increasing unfamiliarity = 0.417, 95% CI = [0.246, 0.588], z = 4.786, p < .001). Therefore, people’s explicit judgments about the structure of the network are behaviorally consistent with the notion that (1) their ability to acquire information about all the friendships in the network is limited, and (2) they use multi-step abstraction to make inferences about potential relationships never observed.

### Abstract cognitive maps of network ties independently encode concurrent relations

To compare different formats of network representation that could potentially support these behavioral patterns, we turn to computational modelling, which allows us to directly characterize the underlying cognitive representations. We consider four possible representational models of the social network. First, as a baseline for comparison, we consider an unlikely (null) model where people’s representations of the network comprise perfect knowledge about all pairwise mutual friendships, such that errors in judgments simply result from noisy application of this knowledge. A second model tests our first main hypothesis that people’s ability to observe friendships is limited by where they sit in the network. This model produces a veridical representation of observations limited by network position, assuming that people are most likely to observe social interactions occurring within their immediate ‘social circle’ and are less likely to observe more distal relations ^45^. We estimate a parameter ω ∈ [0,1] to quantify how strongly observations are affected by this distance: When ω → 0, an individual is estimated to possess only direct information about their own friendships, and as ω → 1, individuals are estimated to have access to information about *all* friendships in the network. To test our second main hypothesis that people abstract over these limited observations, we considered two additional candidate models of multi-step abstraction based on prior work^1–3^: Katz communicability and the Successor Representation (SR). Aside from their distinctive assumptions about the precise representational format of relational information, both models similarly generate abstract representations of associations between every pair of network members as a weighted sum of all the multi-step relations between them, weighing each multi-step relation proportionally based on the number of steps it comprises by a factor of γ ∈ [0,1) ^20,30–32,35,39–41,44^. A direct relation is weighted by γ, while a two-step relation is weighted by γ^2^, etc. When γ → 0, the model reflects direct observations of pairwise social relations. As γ → 1, the representation becomes increasingly abstract, reflecting greater integration of the longer-range connections between a pair of individuals.

At first blush, computational modelling results suggest that, among the candidate models, the Katz model best captures participants’ behavior (average Pearson correlation between model-predicted and observed judgments: mean r = 0.857, 95% CI = [0.815, 0.898], t(99) = 40.857, p < .001; Figure 2a). Its performance is closely followed by the limited observation model (mean r = 0.841, 95% CI = [0.795, 0.885], t(99) = 36.875, p < .001) and then the SR model (mean r = 0.834, 95% CI = [0.790, 0.879], t(99) = 37.218, p < .001). All three models handily outperform the perfect observation (null) model which trails behind in its ability to capture empirical data (mean r = 0.627, 95% CI = [0.590, 0.664], t(99) = 33.362, p < .001). This is confirmed by formal model comparisons that consistently identify Katz as the best model of participants’ behavior (average participant-level model weight based on WAIC: w(WAIC) = 0.605; pxp > .999; overall sample w(WAIC) > .999; Fig 2B). In other words, our results strongly suggest that people’s representations of their social network result from multi-step abstraction over limited observations. Moreover, the fact that Katz also outperforms SR in explaining participants’ inferences about relationships confirms our *a priori* predictions that people’s representations of their social network are structured by default to encode each dyadic relation independently, reflecting the demands of flexible social navigation which includes the ability to track the dynamics of simultaneous social transmission.

**Fig 2.**
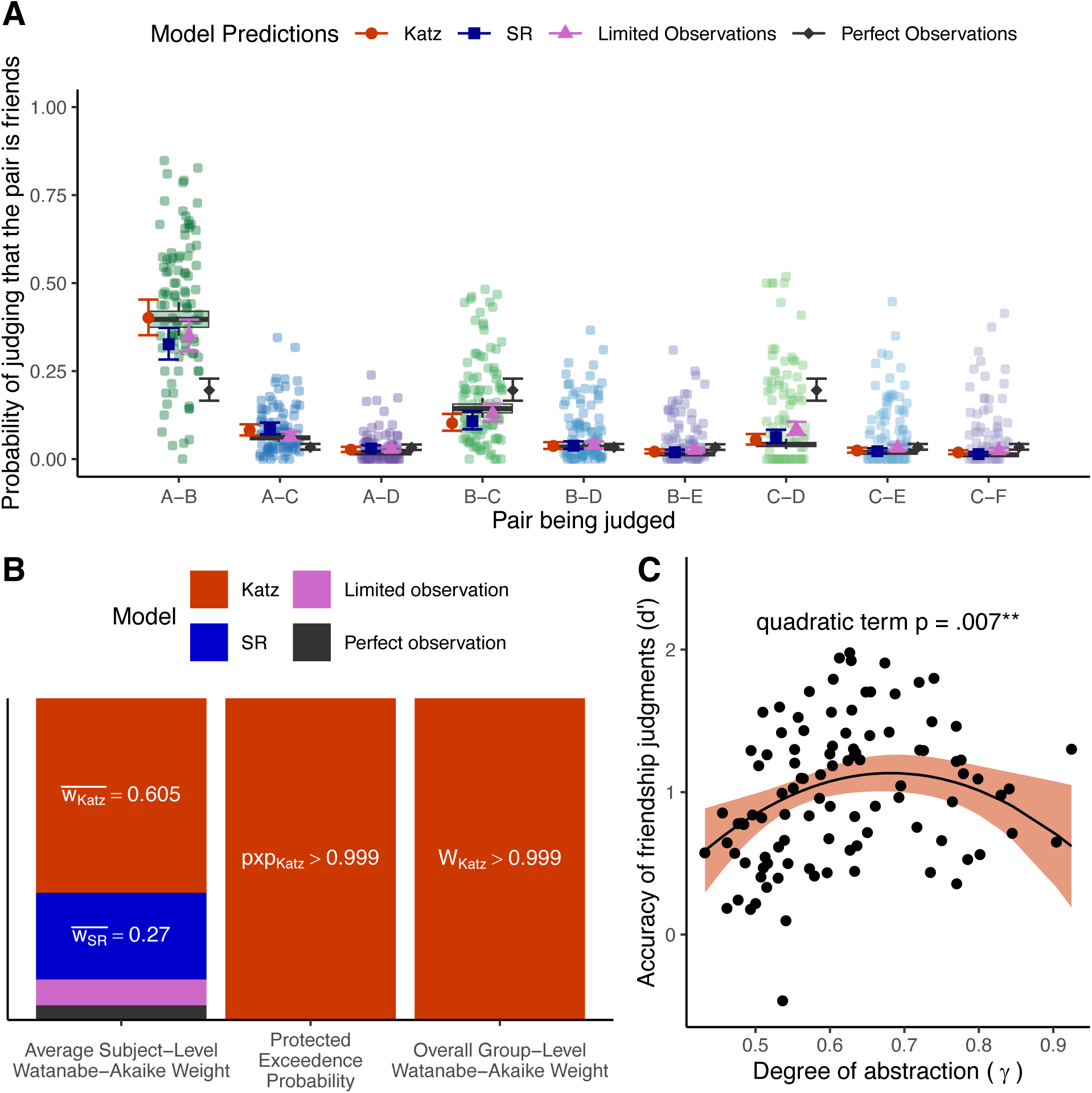
Computational models of network representation. **A) Model fits**. Points and error bars projected over scatterplots and boxplots of participant data (also in Fig 1) indicate the estimated mean and 95% confidence interval based on the within-participant standard error of model predictions averaged across participants. Shape and color of the point and error bars indicate whether the predictions come from Katz, SR, or the models that veridically represent Limited or Perfect Observations. **B) Model comparison**. Height of stacked bars indicate the relative evidence for each of the four candidate models based on the average participant-level model weight based on the Watanabe-Akaike information criterion, protected exceedance probability (pxp) across participants, and the overall group-level Watanabe-Akaike weight. Labels are omitted when model indices < 0.1 (mean w(limited) = 0.080, mean w(perfect) = 0.046). **C) The benefits of abstraction.** The curvilinear relationship between the estimated degree of abstraction (γ) from the best-fitting Katz model and accuracy in the friendship judgment task (d-prime) suggests an optimal degree of abstraction for correctly identifying friendships in the network. Each point indicates a participant. The line and shaded region indicate the estimated mean and 95% confidence interval of predictions from a linear model permitting a quadratic relationship between abstraction and accuracy. **p<.01

To further demonstrate the adaptive utility of abstraction in generating accurate inferences when direct observation is limited, even if it occasionally leads to errors, we examine whether people who engage in more abstraction exhibit more accurate knowledge of friendships in their network. In the best-fitting Katz model, there are two parameters that dictate the accuracy of friendship judgments: being able to directly observe far-away social relationships (ω), and the amount of abstraction over these direct observations (γ). Unsurprisingly, participants estimated to have higher ω were more accurate in their friendship judgments (β = 0.549, 95% CI = [0.072, 1.025], t(95) = 2.286, p = 0.024). Critically, over and above the influence of direct observation, accuracy in friendship judgments was also strongly associated how much a participant abstracts over those observations (γ). This relationship exhibits an inverse u-shape pattern (quadratic term: β = −8.723, 95% CI = [−14.957, −2.489], t(95) = −2.778, p = 0.007; Fig 2C; Supplementary Table S2), such that a moderate level of abstraction yields the greatest benefits for accurately identifying friendships (vertex of the parabola: γ = 0.682), and engaging in any more or less abstraction than this optimum yields lower accuracy. These modelling results support our key hypothesis that people are limited in their ability to construct a high-fidelity representation of their social network, but they can ‘fill in the gaps’ using an appropriate multi-step abstraction mechanism to infer the existence of unobserved social ties.

### Entorhinal cortex encodes abstract cognitive map of one’s social network

A fundamental benefit of constructing abstract cognitive maps of our social networks is that they can be encoded in memory during learning and efficiently retrieved when needed for subsequent choices^29,30^. However, it remains an open question where in the brain these cognitive maps of social networks are stably encoded. We recruited a subset of participants who completed the Network Knowledge Task to return for a separate functional magnetic resonance imaging session (fMRI; N = 43). Participants were presented with photographs of network members, one at a time, and asked to press a button whenever an upside-down image (i.e., a stock photo of a stranger who was not a member of the network) was presented, allowing us to examine whether the spontaneous neural activation of one’s peers naturally comprises information about their social relations in the network^14,46–48^. Using Representational Similarity Analysis (RSA), we estimated how similarly (or dissimilarly) the brain encodes each network member based on the cross-validated Mahalanobis distance between patterns of neural activity ^49,50^. We computed neural similarity in six regions of interest (ROIs) across the medial temporal lobe based on their known role in encoding cognitive maps: left and right anterior hippocampus (HC), left and right posterior hippocampus, and left and right entorhinal cortex (EC; see Methods). We then examined whether (and how strongly) these regions encode participants’ model-estimated (Katz) representation of the network. Each participant’s Katz representation comprises a unique matrix of communicability values reflecting their perception of the overall connectivity between each network member (row) and all other network members (column). Each value reflects the integrated sum of the participant’s direct knowledge of the dyad’s pairwise friendship and their inferences about the dyad’s multi-step connections, controlled by the participant-specific estimates of ω and γ respectively. We first tested whether greater connectivity between two network members—expressed by higher Katz communicability values—predicts more similar patterns of activity in the HC and/or EC (Fig 3A, Supplementary Table S3).

**Fig 3.**
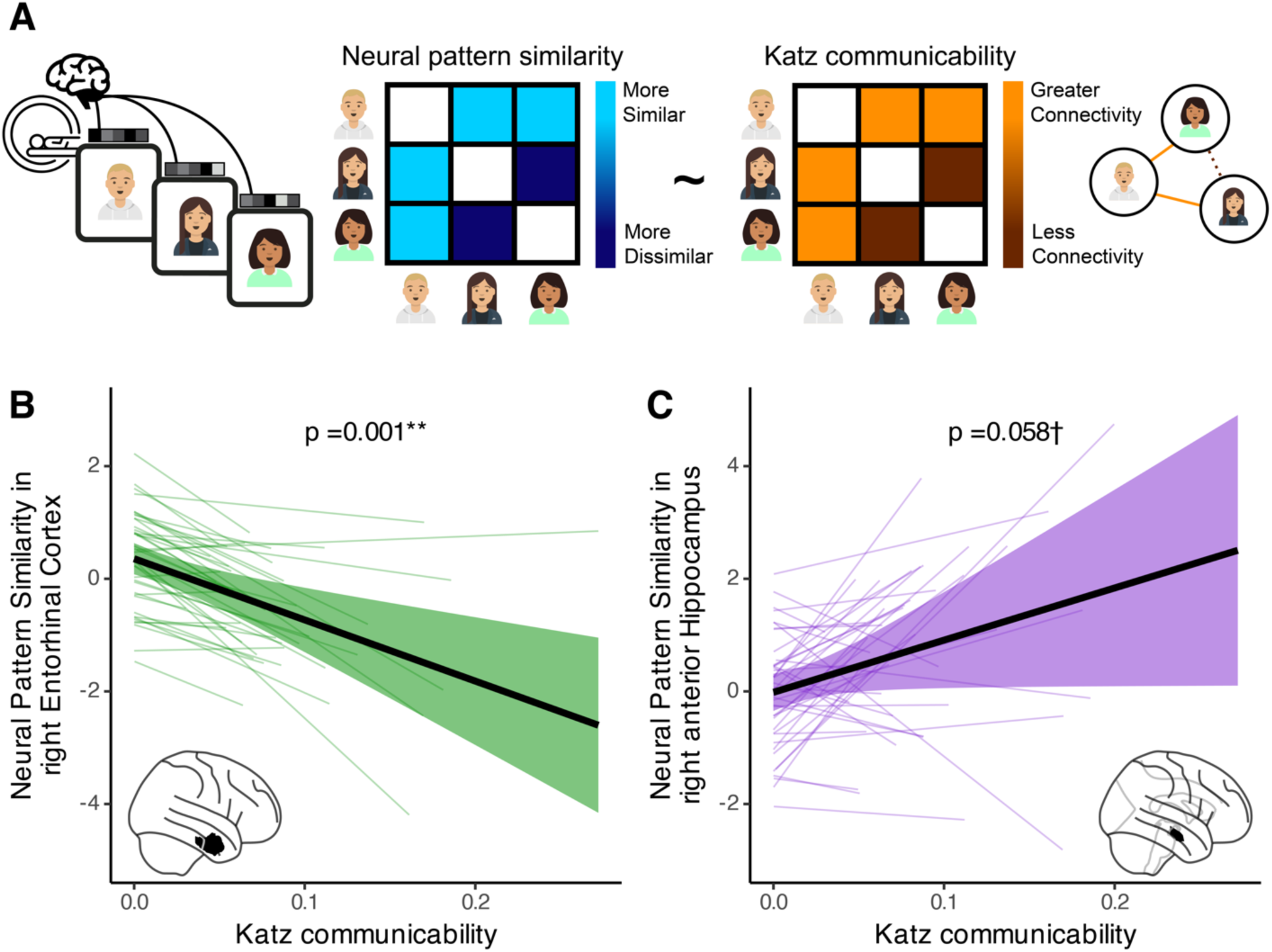
Neural representations of relational maps in a social network. **A) Representational similarity analysis.** To test whether the brain encodes an abstract representation of social network structure, we use Katz communicability between network members to predict neural patterns computed from activity evoked during passive viewing of network members’ photographs during fMRI. Icons: Flaticon.com & getavataaars.com **B) Right entorhinal cortex (rEC) pattern similarity.** Neural pattern dissimilarity in rEC is significantly predicted by Katz communicability. **C) Right anterior hippocampus (raHC) pattern similarity.** Neural pattern similarity in raHC is marginally predicted by Katz. Each thin colored line in B-C indicates the best-fit line for each individual participant from a mixed-effects linear model fitted hierarchically to the data. Solid black lines and colored shaded region indicates the mean and 95% confidence interval of model estimated predictions for the group, respectively. †p < .1, **p < .01 (uncorrected).

We find that the right EC (rEC) encodes network connectivity. The more connected a pair of network members are perceived to be across multi-step relations—i.e. have greater Katz communicability between them—the more they evoked *dissimilar* patterns of neural activity in rEC (rEC ∼ Katz β = −1.087, 95% CI = [−1.706, −0.468], t(31.635) = −3.577, p = 0.001 uncorrected, Bonferroni-corrected at α = .05 for 6 ROIs, p < 0.008; Fig 3B). Although this *dissimilarity* code may seem counterintuitive at first glance, the fact that rEC encodes relevant information about multi-step connectivity illustrates its utility as a functional map of the social network’s structure, which we explore more fully in subsequent analyses. In the right anterior HC, we only observed marginal effects of connectivity between network members on neural pattern similarity (raHC ∼ Katz β = 0.927, 95% CI = [−0.035, 1.888], t(29.314) = 1.970, p = 0.058 uncorrected; Fig 3C). There was no evidence that relational information is encoded in the right posterior HC or any of the left-lateralized ROIs (Ps ≥ .255), or the control visual regions (left and right V1: Ps ≥ .225).

To unpack these neural results, we conducted two additional follow-up analyses to elaborate on the specific nature and content of the observed representations. Our results suggest that the rEC deploys a dissimilarity code to differentiate network members who exhibit greater connectivity. Although the contributions of EC to pattern separation and differentiation in the HC are well-documented^51–53^, only recently has evidence emerged that the EC itself engages in pattern separation and differentiation^54^. We therefore wanted to understand if the rEC uses a dissimilarity code to resolve potential interference^55–58^. In this case, the natural prediction is that the brain maximizes the distinctiveness of representations for network members closest to the participant, where misidentification about nuanced properties of the individual (e.g., what are their likes, dislikes, etc.) likely incurs the greatest social costs. To do this, we examined whether rEC pattern similarity between network members changes as a function of participants’ familiarity with the dyad. rEC patterns are most dissimilar when participants are directly friends with at least one member of the dyad, and these patterns of neural activity become less dissimilar as participants become more distant to the dyad (unfamiliarity β = 0.224, 95% CI = [0.019, 0.429], t(41.234) = 2.208, p = 0.033, Supplementary Table S4). However, this alone was not sufficient to explain the content of the rEC neural code. Simultaneous inclusion of unfamiliarity *and* Katz communicability in a regression model instead confirms that multi-step connectivity robustly predicts pattern differentiation in rEC (Katz β = −0.856, 95% CI = [−1.477, −0.234], t(34.278) = −2.797, p = 0.008, Supplementary Table S4). These analyses strongly suggest that the dissimilarity code in rEC reflects its fundamental role in preventing interference between representations that share many features^51,51,52,54^—i.e., multi-step connections in our case of social networks.

Second, because model-estimated Katz communicability values reflect the integration of both inferred multi-step relations and direct observations of pairwise ties, it is unclear whether neural encoding of a Katz representation merely reflects the memorization of observed ties, or if these neural processes integrate and encode inferences of unobserved relations through multi-step abstraction. To probe whether the abstracted inferences are indeed encoded in the brain, we conducted follow-up analyses that control for the possible observations the participant could make within our computational model (Supplementary Table S5). We find that Katz-encoding in the rEC remains significant (b_Katz_ = −1.495, 95% CI = [−2.408, −0.581], t(38.320) = −3.312, p = 0.002), while the marginal association between Katz and neural pattern similarity in raHC is abolished (b_Katz_ = 0.924, 95% CI = [−0.958, 2.807], t(16.390) = 1.039, p = 0.314). In combination with additional supplementary analyses (Supplementary Table S6-S7), these findings establish the key role of rEC in robustly encoding an abstracted map of multi-step connections within one’s social network, while also suggesting that the raHC might be encoding a more veridical representation of observed social ties.

### Neural encoding of relational maps for tracking information flow

Although the rEC appears to encode relational maps of one’s social network, what is the functional utility of these maps for solving navigation problems in the network? One formidable social navigation problem is understanding how information flows across chains of social ties. If an individual understands how information might traverse across the network, it enables several adaptive behaviors, from strategically spreading gossip that can sabotage relationships without getting caught, to enhancing one’s reputation by acting generously towards well-connected members. Successfully deploying these strategic behaviors requires an intricate understanding of how the litany of social ties enable information to spread. To test whether socio-cognitive maps in rEC and raHC facilitate predictions about information flow, we probed participants’ inferences about information flow in a separate task conducted independently from the fMRI session (see Methods). In this Information Flow Task, participants were instructed that a network member (the source) has hypothetically disclosed some news about themselves, and then asked to judge how likely it would be for another network member (the target) to hear this news. Accurate inference in this task requires knowledge of whether, and how, two individuals are connected, and ideally, knowledge of every possible connection between them—as news can travel on any number of paths. Given this, we tested the hypothesis that neural maps tracking multi-step connectivity between network members would be valuable for tracking information flow.

Consistent with this prediction (and our finding that Katz communicability is represented using a *dissimilarity* code in rEC), greater dissimilarity between rEC activity evoked by the source and target is associated with inferences that information is more likely to flow between them (β = −0.273, 95% CI = [−0.434, −0.113], z = −3.336, p < .001; Fig 4A-B, Supplementary Table S8). Additionally, consistent with our results suggesting that Katz communicability may be represented using a *similarity* code in raHC, greater similarity between neural patterns evoked by the source and target in the raHC is also associated with inferences that information flows between them (β = 0.157, 95% CI = [0.017, 0.296], z = 2.197, p = 0.028; Fig 4C). Furthermore, because these analyses consider the neural patterns from both these regions simultaneously, our findings suggest that neural patterns in rEC and raHC are independently and uniquely associated with judgments of information flow. Put differently, when trying to infer whether information would spread from one person to another in their social network, people draw on both the relational maps encoded in the rEC and raHC.

**Fig 4.**
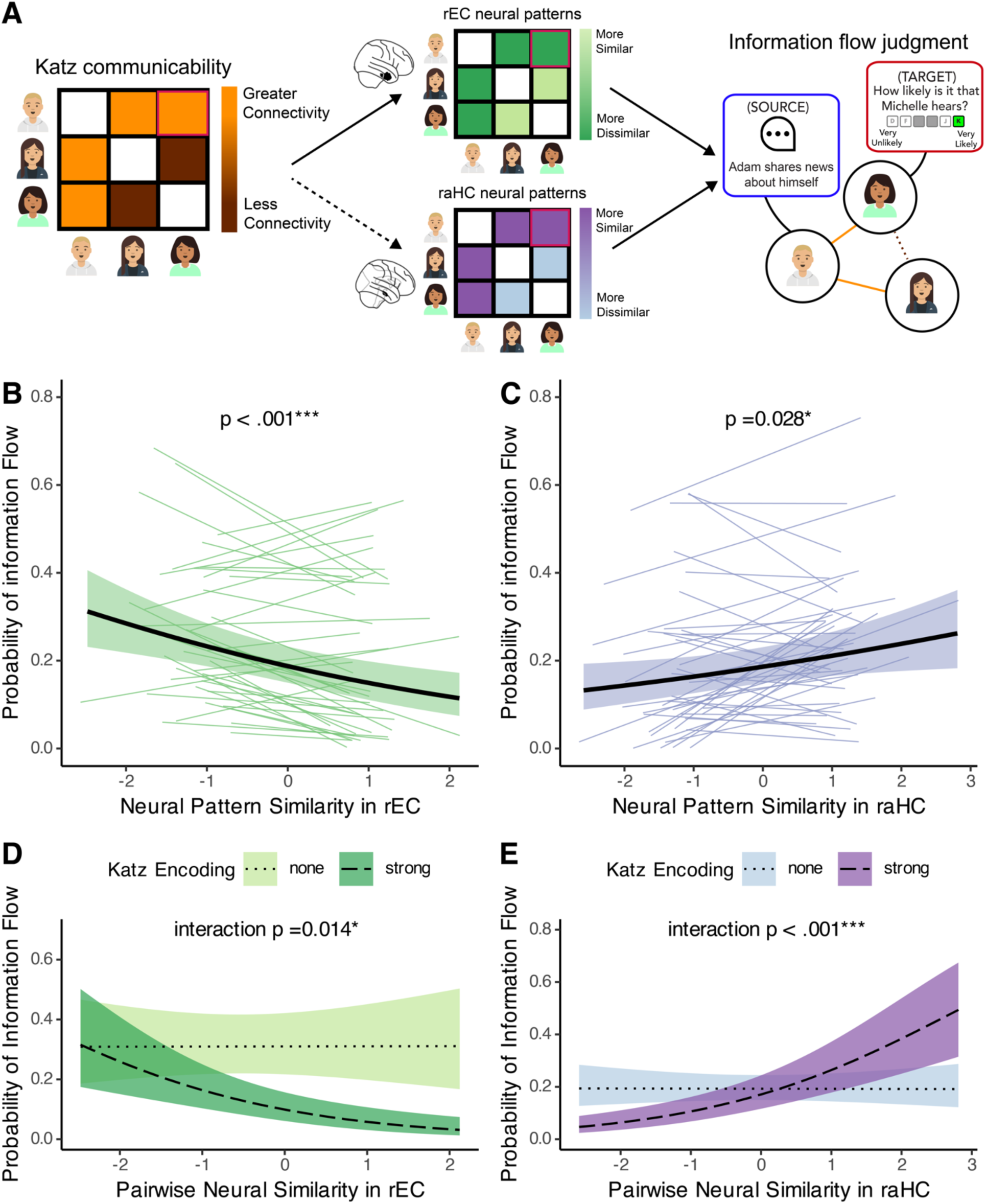
Neural representation of abstract relational maps supports flexible social decision-making. **A) Predicted association between neural representations and information flow judgment.** Because Katz communicability is encoded as greater pattern dissimilarity in rEC and greater similarity in raHC, inferences of information flow are supported by pattern dissimilarity in rEC and pattern similarity in raHC, we expect information flow judgments to be supported by pattern dissimilarity in rEC and pattern similarity in raHC. Icons: getavataaars.com **B) Group-level effect in rEC.** As Katz communicability is encoded using a dissimilarity code in rEC, inference of information flow is supported by pattern dissimilarity. **C) Group-level effect in raHC.** As Katz may rely on a similarity code in raHC, inferring information flow is supported by pattern similarity. Each thin line in B-C indicates the best-fit line for each individual participant from a mixed-effects linear model fitted hierarchically to the data. Solid black lines and shaded regions in B-C indicates the mean and 95% confidence interval of the average model estimated predictions respectively for the group. **D) Individual differences in rEC.** The more strongly an individual’s rEC encodes Katz communicability, the more strongly rEC predicts inference of information flow. **E) Individual differences in raHC.** The more strongly an individual’s raHC encodes Katz communicability, the more strongly raHC predicts inference of information flow. Lines and shaded regions in D-E indicate the mean and 95% confidence interval of model-estimated probabilities of information flow between two network members, as a function of neural pattern similarity and level of Katz encoding in the respective regions. Dotted lines and light shading indicate the model-estimated probabilities when participants’ neural patterns do *not* encode Katz (none: neural patterns ∼ Katz β = 0). In contrast, dashed lines and dark shading indicate model estimates when participants exhibit strong levels of Katz encoding (strong: rEC patterns tracks very negatively with Katz, mean −1SD: β = −2.296; raHC patterns tracks very positively with Katz, mean + 1SD: β = 3.234). *p < .05, ***p < .001.

**Fig 5.**
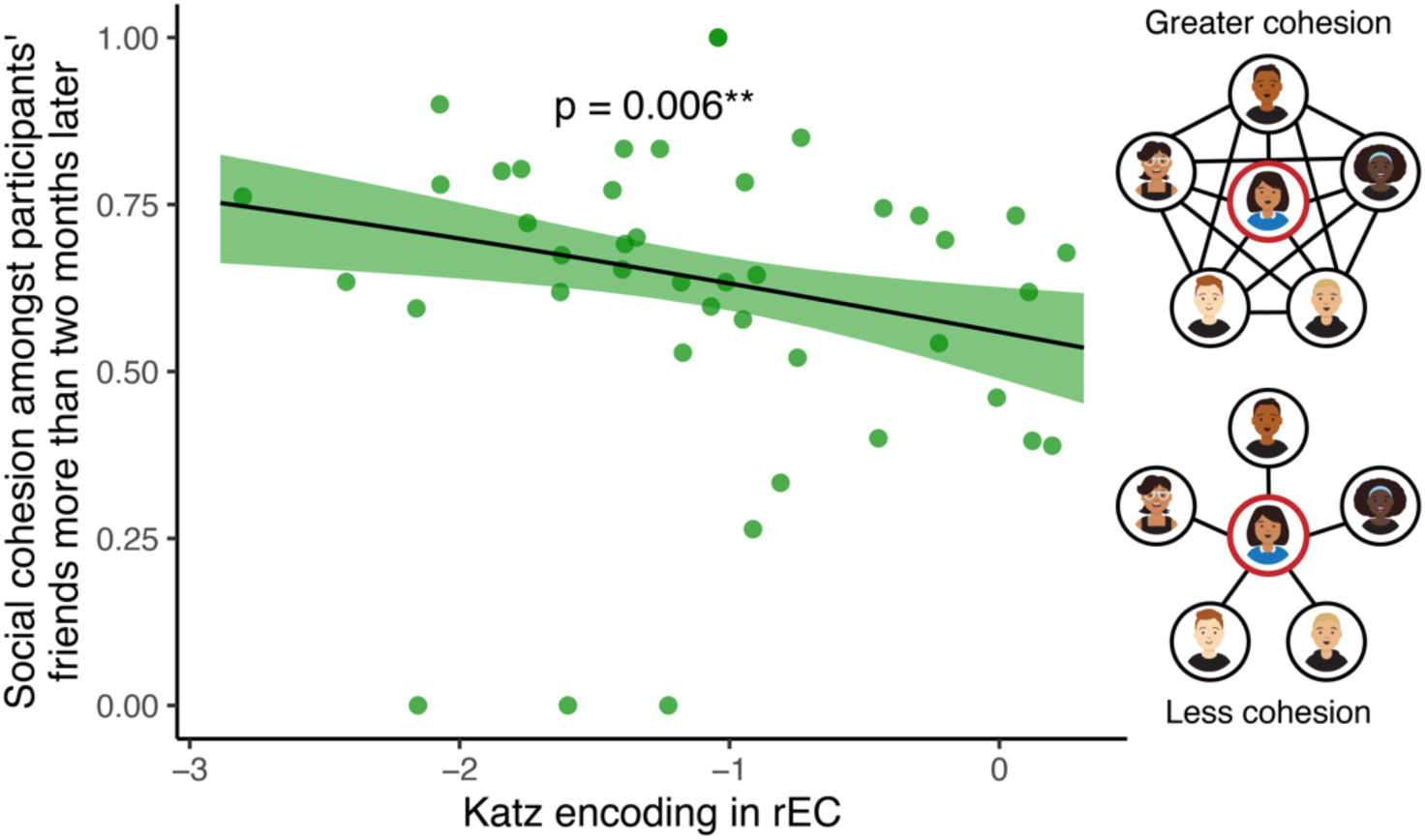
Abstract maps of multi-step relations in rEC afford effective social brokerage. The degree to which an individual encodes Katz communicability in the rEC predicts greater social cohesion amongst their friends (i.e., more effective brokerage) more than two months later. Each point represents a single participant. Because the rEC represents the multi-step connectivity between dyads in the network using a dissimilarity code, negative values of Katz encoding (x-axis) indicate greater encoding strength. The black line and shaded region indicate the predicted mean and 95% confidence interval from an ordered beta regression model. Having controlled for social cohesion amongst friends when neural activity was measured, this association can be effectively interpreted as the relative change in social cohesion over time. **p < .01. Icons: getavataaars.com

If the rEC and raHC support tracking of information flow primarily by encoding efficient maps of the social network for subsequent retrieval, then we might expect that failures to encode these maps render the neural codes in these regions uninformative for navigational inference. To test this, we leverage the considerable individual differences in how strongly participants encode a Katz representation in rEC and raHC. We then ask whether inferences in the Information Flow Task are better predicted by neural pattern similarity in people who encoded Katz more strongly in these regions. As predicted, we find that our ability to use neural pattern (dis)similarity to predict a participant’s behavioral inferences about information flow depends on how strongly participants encode a Katz representation in our regions of interest (modulation of rEC patterns: β = 0.253, 95% CI = [0.052, 0.455], z = 2.461, p = 0.014; modulation of raHC patterns: β = 0.172, 95% CI = [0.099, 0.245], z = 4.630, p < .001; Fig 4D-E).

In other words, for someone whose rEC neural patterns strongly differentiate network members based on their Katz communicability (i.e., more negative Katz encoding) these neural codes reliably predict their inferences about whether information spreads from one member to another (simple effect β = −0.579, 95% CI = [−0.886, −0.273], z = −3.704, p < .001 at Katz encoding = mean −1SD). For someone whose rEC neural patterns are unrelated to Katz and thus contain no information about connectivity (i.e., Katz encoding = 0) rEC pattern dissimilarity for a dyad is unrelated to participants’ inferences about whether information flows between its members (simple effect β = 0.002, 95% CI = [−0.255, 0.259], z = 0.016, p = .987).

Similarly, for someone whose raHC neural representations exhibit greater similarity for network members who have high Katz communicability (i.e., more positive Katz encoding), these codes reliably predict their inferences about whether information spreads from one member of a dyad to the other (simple effect β = 0.553, 95% CI = [0.344, 0.763], z = 5.178, p < .001 at Katz encoding = mean+1SD). For someone whose raHC are unrelated to Katz and thus contain no information about connectivity (i.e., Katz encoding = 0) raHC pattern similarity between members of the dyad is again unrelated to participants’ inferences about how information flows (simple effect β = −0.003, 95% CI = [−0.163, 0.157], z = −0.035, p = .972). Together, these results provide strong evidence that the rEC and raHC’s encoding of multi-step connections within the social network is consequential for tracking information flow.

### rEC maps of the social network enable effective social brokerage

There are of course other types of social navigation problems that go beyond inferences about information flow. Another notable feature of social networks is that each member and their social ties actively contributes to its evolution over time. Thus, we might also ask whether abstract neural maps of multi-step relations afford individuals a unique ability to shape the social communities they are a part of. As we have already illustrated, possessing an abstract neural map enables the identification of multi-step connectivity between people. Naturally, this also includes the connectivity amongst one’s friends. With the spontaneous availability of this knowledge in their daily lives, one might then be able to subsequently broker the formation of friendships between friends who are currently unconnected to each other to foster a more cohesive social community, which ultimately affords the individual greater social capital^1^. To test this, we recontacted 176 participants from our initial network (N = 187) in the following semester at college, two months after neuroimaging, to identify their friendships again. This allows us to investigate whether the encoding of abstract neural maps earlier in the year is associated with greater social cohesion amongst participants’ friends more than two months later.

Within a social network, the cohesion amongst an individual’s friends can be quantified by (the inverse of) the average number of connections separating each pair of their friends if the participant were to be removed from the network—a metric known as local efficiency ^59,60^. In other words, greater distances between a participant’s friends if the participant is removed from the network indicates poor cohesion amongst the group. In contrast, a maximally cohesive social group would comprise individuals that are all mutually friends with each other. Confirming our hypothesis, we find that participants who more strongly encode an abstract neural map in the rEC (i.e., more negative Katz encoding) have more cohesive friend groups later in the academic year (β = −0.303, 95% CI = [−0.519, −0.086], z = −2.737, p = 0.006). This was not the case for raHc encoding (β = 0.084, 95% CI = [−0.017, 0.185], z = 1.638, p = 0.101), or the control regions of V1 (Ps>.3, Supplementary Table S9). Having controlled for the social cohesion of their friend group prior to the neuroimaging session used for this analysis (see Table 1), these associations imply that the availability of the rEC neural maps facilitates *increases* in the cohesion of one’s social group: people who strongly encode these neural maps of their social network possess more cohesive friend groups independent of initial levels of cohesion. Specification of the model to control for social cohesion measured *after* neuroimaging as an alternative baseline confirms that the availability of neural maps robustly *predicts* increases in the social cohesion of their communities rather than simply reflect changes in the network between the initial survey and scanning (Supplementary Table S10). Moreover, further analyses demonstrate that these increases are neither driven by (1) participants leaving their communities and moving into other highly cohesive communities (Supplementary Table S11), nor (2) the relative degree of abstraction people engage in (Supplementary Table S12). Rather, the spontaneous availability of an abstract neural map of people’s network in the rEC specifically seems to afford them the ability to draw on the kinds of relational information necessary to broker the formation of friendships between their friends, fostering greater social cohesion in their immediate communities over time.

**Table 1:**
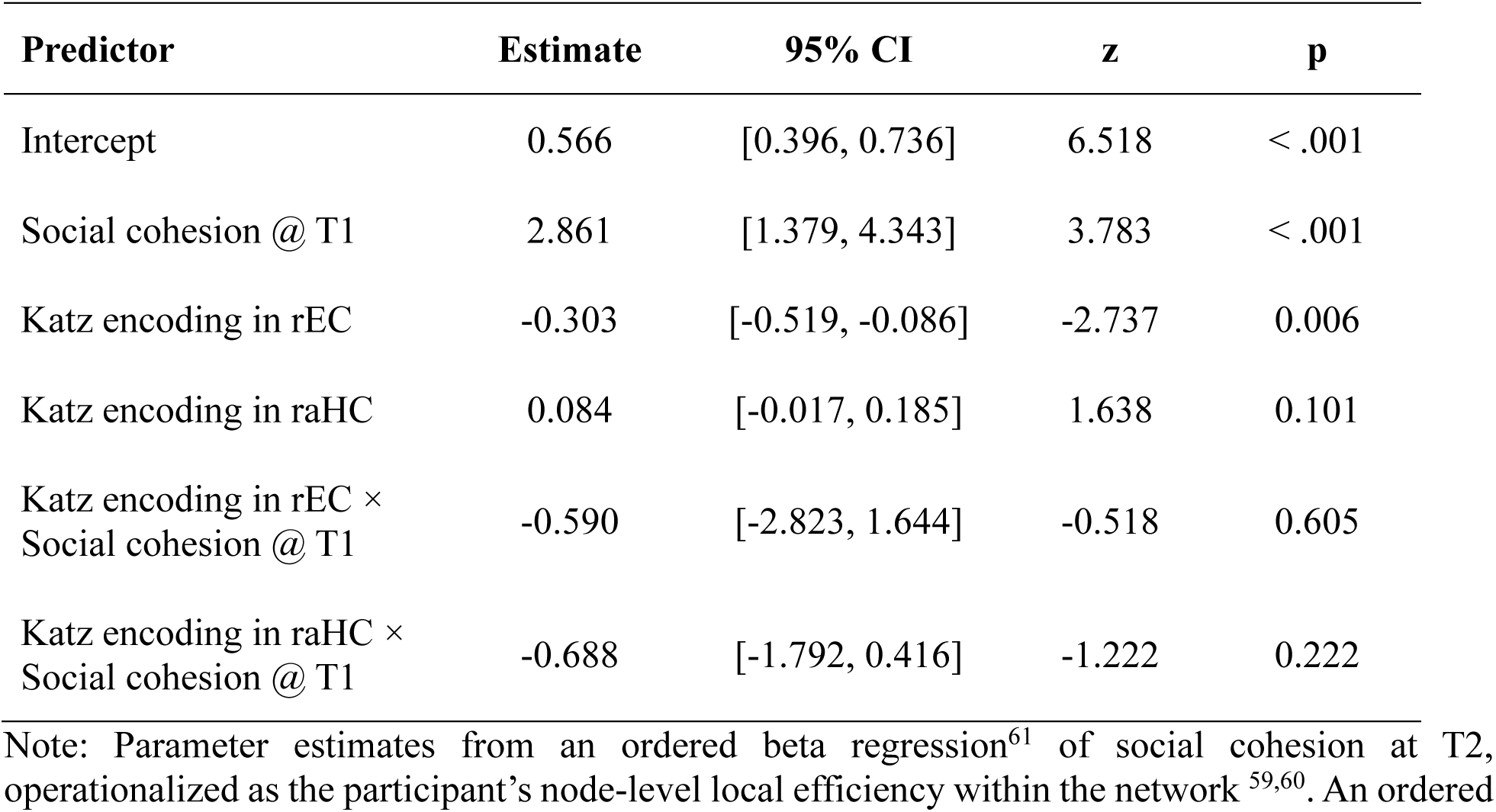
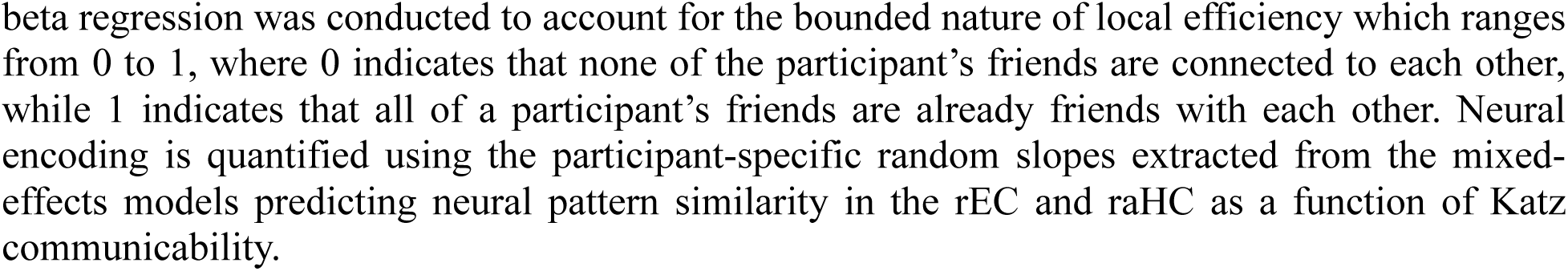
Association between neural encoding of abstract relational maps and future social cohesion.

Note: Parameter estimates from an ordered beta regression^61^ of social cohesion at T2, operationalized as the participant’s node-level local efficiency within the network ^59,60^. An ordered beta regression was conducted to account for the bounded nature of local efficiency which ranges from 0 to 1, where 0 indicates that none of the participant’s friends are connected to each other, while 1 indicates that all of a participant’s friends are already friends with each other. Neural encoding is quantified using the participant-specific random slopes extracted from the mixed-effects models predicting neural pattern similarity in the rEC and raHC as a function of Katz communicability.

## Discussion

People are remarkably proficient at navigating a wide array of social situations that require them to keep track of the intricate web of relations between members of a social network ^31,32,35,62^. However, it has remained unknown exactly how the human brain organizes such forms of relational information into a useful representation for flexible behavior in the real-world, especially when networks are large and complex and direct observations about specific relationships are scarce. We combined computational modelling, functional neuroimaging, and longitudinal social network analysis of a large and emerging real-world network to answer this question in a naturalistic and ecologically valid setting. Despite constraints on being able to directly acquire knowledge about all other friendships in one’s community ^31,33^, we find that people correctly identify friendships in their personal social network far above chance levels. Consistent with recent work^31,32^, people accomplish this feat by constructing abstract representations of their social networks which integrate not only direct knowledge of friendships, but also predictive inferences about unobserved (but likely) relations between people.

Our results also reveal the precise format of these representations. We find that the Katz model^35,40^ which prioritizes the independent—rather than relative—encoding of people’s concurrent relationships^38,39,41^ handily outperforms the Successor Representation in explaining our participants’ behavior^31,32^. Prior work using laboratory tasks has reported evidence for people employing an SR in non-social^18,30,43,44^ and in social domains^31,32^. Our results suggest that representations of real-world social environments are naturally structured to capture the specific dynamics of everyday social phenomena, such as how information simultaneously flows across multiple chains of relationships

Furthermore, we provide direct evidence that a Katz-based abstract map of the social network is encoded in regions of the medial temporal lobe: namely, the right entorhinal cortex (rEC), and to a lesser extent, the right anterior hippocampus (raHC). Extant work has established that these same regions are implicated in constructing and maintaining map-like representations of relational information^8,10,12,15,17–28^, including representations that resemble an SR in other domains^18,20^. Our finding that real-world social network structure is encoded in the MTL adds to this literature and argues for domain-general mapping functions of this region. On the other hand, our finding that these representations encode Katz communicability illuminates how common neural mechanisms of abstraction might adapt the precise format of these representations to specific properties of the relational structures they encode, which in turn enables flexible and adaptive inferences across different environments that range from simple grid-worlds to complex non-Euclidean social networks.

Furthermore, we demonstrate the versatility of these neural representations for adaptive social behavior. We find that these abstract neural codes support people’s inferences about how information spreads across their social network (assessed in a completely separate and independent task). This ability to track the flow of information through social ties within a vast network provides a potential mechanistic explanation for how people evaluate the reputational effects of their social behavior and flexibly adapt their actions to actively manage these considerations that govern social life^4,35,42,63–66^. Simultaneously, we show that these neural codes appear to afford individuals the ability to meaningfully shape the evolution of their immediate social communities. By tracking the multi-step connectivity between one’s own friends, these maps identify opportunities for people to broker connections between unconnected individuals within the group and foster greater social cohesion. While our work cannot directly show that people deploy these maps to bring their communities together, we find that it is the *spontaneous availability* of these neural maps in a passive viewing task that predicts future increases in social cohesion of one’s social group. As a result, these effects do not simply indicate that people who infer strong connectivity between peers who share multi-step relations (e.g., my friends who are connected through me) accurately anticipate that they are likely to become friends. Instead, it is the degree to which these inferences are spontaneously activated during people’s interactions with others that predict increases in the cohesion of their social community. We speculate that this is because spontaneous inferences about multi-step connectivity between one’s friends in daily life might promote actions to bring them into contact with one another, thereby brokering a direct friendship between those who are currently unconnected. Critically, given the socio-psychological benefits of inhabiting a highly cohesive social community for well-being^67,68^, our results suggest that possessing these neural maps might uniquely endow individuals with a powerful form of social capital^1,69^.

Our findings lay the important foundation towards developing a more comprehensive understanding of how the human brain supports the ability to skillfully navigate our complex real-world social networks across a wide range of situations. By showing that rEC engages in pattern differentiation, our findings add to the emerging evidence that EC plays a direct role in interference resolution between memories that share similar features^54^ and not simply through its inputs into the hippocampus^51–53,55,58,70^. We suspect that the deployment of this dissimilarity code in the EC reflects the utility of preventing interference between distinct individuals who occupy similar structural positions within the social network—i.e., those who share a great deal of mutual and multi-step connections^71,72^. Future work should more thoroughly interrogate this hypothesis.

Another potential avenue for future work concerns the preliminary evidence that neural representations of network relations in the rEC and raHC may be attuned to distinct components of network knowledge: the rEC seems to preferentially encode an inferential map of multi-step connections while the raHC seems to preferentially encodes a more veridical map of relationships based on directly acquired information—though this latter claim is qualified by weaker evidence. A potential source of uncertainty around the raHC representation comes from the fact that we have limited insight into both what (and how much) direct information people are able to acquire about others’ social relationships ‘in the wild.’. This requires that we estimate what people might be observing based on where they sit in the network ^31,45^. In reality, the relational information directly available to people is likely more diverse and idiosyncratic, and future work would have to develop more sophisticated techniques to circumvent the challenges of measuring these observations in real-world networks to confirm whether the brain separately encodes veridical memory for observed relationships.

Setting this aside, our work touches on past theories that medial temporal lobe representations might exhibit a potential gradient of abstraction, with more abstract representations encoded in the EC^15,17^ compared to the HC ^12,18–20^. Such forms of complementary neural encoding might afford greater cognitive flexibility across a range of social decisions ^16^. Selective use of more abstract representations in EC can be used when generalizing about unobserved or unknown relationships, and more veridical HC representations can be relied upon when greater precision is required. Future work investigating how people incrementally construct and update these integrated neural network representations from discrete, piecemeal information about pairwise relationships, and how these representations in the EC and HC are differentially re-instantiated during choice will help shed light on these open questions.

## Methods

### Overview

We recruited 198 first-year undergraduate students across three dormitories within the first month of their arrival on campus (N = 196 after 2 exclusions for not meeting eligibility; collected Sep 2022; see Table 2 for demographic breakdown). At this initial time point, they completed demographic questionnaires, submitted photographs of themselves to be used as stimuli for subsequent data collection, and provided informed consent to be contacted for subsequent waves of data collection. Participants were then recontacted six times throughout the academic year to complete a survey in which they identified their friends amongst the 195 other participants. They were also recontacted another three times to complete the Network Knowledge Task and Information Flow Task, and once to participate in a functional neuroimaging session.

**Table 2:** Network Demographics.

|  | Initial<br>Recruitment<br>(N =196) | Friendship<br>Survey 2<br>(N = 187) | Network<br>Knowledge &<br>Information<br>Flow<br>(N = 100) | Functional<br>Neuroimaging<br>(N = 43) | Friendship<br>Survey 3<br>(N = 180) | Friendship<br>Survey 4<br>(N = 176) |
| --- | --- | --- | --- | --- | --- | --- |
| **Gender** |  |  |  |  |  |  |
| Female | 103 | 100 | 54 | 26 | 95 | 96 |
| Male | 89 | 83 | 44 | 17 | 81 | 77 |
| Other | 4 | 4 | 2 | 0 | 4 | 3 |
| **Ethnicity** |  |  |  |  |  |  |
| Asian | 54 | 52 | 27 | 11 | 51 | 50 |
| Black | 23 | 22 | 14 | 5 | 21 | 21 |
| East Asian | 5 | 5 | 3 | 2 | 4 | 5 |
| Hispanic/Latinx | 21 | 21 | 13 | 5 | 21 | 20 |
| Middle Eastern | 4 | 4 | 1 | 1 | 4 | 3 |
| Mixed | 18 | 17 | 8 | 3 | 16 | 16 |
| Native<br>American | 1 | 1 | 0 | 0 | 1 | 0 |
| South Asian | 16 | 14 | 12 | 4 | 14 | 14 |
| White | 50 | 47 | 21 | 12 | 44 | 43 |
| Other | 4 | 4 | 1 | 0 | 4 | 4 |
| **Age** |  |  |  |  |  |  |
| Mean | 18.1 | 18.1 | 18.1 | 18.1 | 18.1 | 18.1 |
| SD | 0.421 | 0.424 | 0.424 | 0.402 | 0.420 | 0.425 |
| Min | 17 | 17 | 17 | 17 | 17 | 17 |
| Max | 19 | 19 | 19 | 19 | 19 | 19 |

Given our goal of identifying and characterizing the neural representations of the social network, we restricted our analyses to data collected during sessions directly adjacent to the functional neuroimaging session (N = 43 after 4 exclusions for data quality; collected Nov 2022). This included the second friendship survey collected approximately one month after the study began (N = 187; collected Oct 2022, T1), and the first sessions of the Network Knowledge Task and Information Flow Task (N = 100; collected Nov 2022 on separate occasions at least one week apart). Finally, to assess how participants’ social communities evolved over time, we examined changes in people’s friendships between the second friendship survey and the fourth friendship survey (N = 176; collected Feb 2023, T2). To ensure that our observed associations were indicative of predictions and did not simply reflect changes in network structure that occurred between the second survey and our functional neuroimaging session, supplementary analyses confirm the robustness of these effects using the third friendship survey as an alternative baseline for comparison instead of the second (third friendship survey; N = 180; collected Dec 2022, T1.5). All procedures were approved by Brown University’s Institutional

Review Board, and informed consent was obtained from all subjects. For 17-year-old minors, informed consent was obtained from their legal guardians, and assent was obtained from the subjects.

### Network Knowledge Task

In this task, participants were presented with the name and photograph of a network member (target) alongside a list of names and photographs of other network members (probes), and asked to judge whether the two were friends. Given the size of the network, we restricted each participant’s sample of targets and probes to a subset of 30 network members based on their network distance to the participant. Where possible, the personalized stimuli set consisted of 5 randomly sampled friends (distance = 1), 10 randomly sampled friends-of-friends (distance = 2), and 15 randomly sampled friends-of-friends-of-friends (distance = 3). If there were insufficient network members at a specific level of network distance for a participant, we added members at the next level of distance to fill out the stimulus set. Each participant made 870 friendship judgments for every pairwise combination of the thirty network members (0: not friends, 1: friends), and rated each pair twice: Each member of a pair appears once as the target and once as the probe.

### Information Flow Task

In this task, participants were presented with the name and photograph of a network member (source) and told that they had disclosed some news about themselves. Participants were then asked how likely another network member (target), whose name and photograph were also presented, would hear about this news on a four-point scale that ranged from ‘very likely’ to ‘very unlikely’. Stimuli for this task comprised a personalized subsample of 25 network members that were also presented in the Network Knowledge Task, and thus similarly varied in their network distance from the participant. Participants made these information flow judgments in blocks of 24 trials, where each block focused on one of the 25 network members as the Source and each trial in a block elicited likelihood judgments for each of the 24 other network members as targets, resulting in a total of 600 judgments Responses from this task were binarized for subsequent analyses (0: very unlikely or unlikely; 1: likely or very likely).

### Functional neuroimaging

#### Task

Participants were scanned while passively viewing photographs of a further subset of 22 network members previously used as stimuli in the Information Flow and Network Knowledge Tasks^46–48^. To ensure participants were visually processing the identity of the network members, participants were instructed to make a button press when an upside-down face (stock image) was presented instead of a real network member. Each network member in the stimulus set and the one upside-down face was presented to participants three times resulting in a total of 69 trials every run. A fixation cross was displayed during the interstimulus intervals (ISIs) for a jitter of 1 to 8 s (mean duration = 3.5 s). Participants completed seven functional runs, each lasting six minutes.

#### MRI acquisition

Data was acquired at the Brown University MRI Research Facility with a Siemens Prisma 3T MRI scanner. Anatomical scans were collected using a standard T1-weighted magnetization prepared rapid acquisition gradient echo (MPRAGE) sequence (1mm^3^ isotropic voxels, TR = 1900ms, TE = 3.02ms, flip angle = 9 degrees, 160 slices). Functional scans were collected using a T2*-weighted echo-planar imaging (EPI) sequence that prioritized coverage of hippocampus and entorhinal cortex (3mm^3^ isotropic voxels, TR = 2000ms, TE = 25ms, flip angle = 90 degrees, 40 slices).

#### fMRIPrep boilerplate

Results included in this manuscript come from preprocessing performed using *fMRIPrep* 21.0.0 (RRID:SCR_016216)^73,74^, which is based on *Nipype* 1.6.1 (RRID:SCR_002502)^75,76^. The below boilerplate text was automatically generated by fMRIPrep with the express intention that users should copy and paste this text into their manuscripts *unchanged*. It is released under the CC0 license.

#### Anatomical data preprocessing

A total of 1 T1-weighted (T1w) images were found within the input BIDS dataset. The T1-weighted (T1w) image was corrected for intensity non-uniformity (INU) with N4BiasFieldCorrection^77^, distributed with ANTs 2.3.3 (RRID:SCR_004757)^78^, and used as T1w-reference throughout the workflow. The T1w-reference was then skull-stripped with a *Nipype* implementation of the antsBrainExtraction.sh workflow (from ANTs), using OASIS30ANTs as target template. Brain tissue segmentation of cerebrospinal fluid (CSF), white-matter (WM) and gray-matter (GM) was performed on the brain-extracted T1w using fast (FSL 6.0.5.1:57b01774, RRID:SCR_002823)^79^. Volume-based spatial normalization to one standard space (MNI152NLin2009cAsym) was performed through nonlinear registration with antsRegistration (ANTs 2.3.3), using brain-extracted versions of both T1w reference and the T1w template. The following template was selected for spatial normalization: *ICBM 152 Nonlinear Asymmetrical template version 2009c* (RRID:SCR_008796; TemplateFlow ID: MNI152NLin2009cAsym)^80^.

#### Functional data preprocessing

For each of the 7 BOLD runs found per subject (across all tasks and sessions), the following preprocessing was performed. First, a reference volume and its skull-stripped version were generated using a custom methodology of *fMRIPrep*. Head-motion parameters with respect to the BOLD reference (transformation matrices, and six corresponding rotation and translation parameters) are estimated before any spatiotemporal filtering using mcflirt (FSL 6.0.5.1:57b01774)^81^. BOLD runs were slice-time corrected to 0.965s (0.5 of slice acquisition range 0s-1.93s) using 3dTshift from AFNI (RRID:SCR_005927)^82^. The BOLD time-series (including slice-timing correction when applied) were resampled onto their original, native space by applying the transforms to correct for head-motion. These resampled BOLD time-series will be referred to as *preprocessed BOLD in original space*, or just *preprocessed BOLD*. The BOLD reference was then co-registered to the T1w reference using mri_coreg (FreeSurfer) followed by flirt (FSL 6.0.5.1:57b01774)^83^ with the boundary-based registration^84^ cost-function. Co-registration was configured with six degrees of freedom. Several confounding time-series were calculated based on the *preprocessed BOLD*: framewise displacement (FD), DVARS and three region-wise global signals. FD was computed using two formulations following Power (absolute sum of relative motions)^85^ and Jenkinson (relative root mean square displacement between affines)^81^. FD and DVARS are calculated for each functional run, both using their implementations in *Nipype* (following the definitions by ^85^). The three global signals are extracted within the CSF, the WM, and the whole-brain masks. Additionally, a set of physiological regressors were extracted to allow for component-based noise correction (*CompCor*)^86^. Principal components are estimated after high-pass filtering the preprocessed BOLD time-series (using a discrete cosine filter with 128s cut-off) for the two *CompCor* variants: temporal (tCompCor) and anatomical (aCompCor). tCompCor components are then calculated from the top 2% variable voxels within the brain mask. For aCompCor, three probabilistic masks (CSF, WM and combined CSF+WM) are generated in anatomical space. The implementation differs from that of Behzadi et al. in that instead of eroding the masks by 2 pixels on BOLD space, the aCompCor masks are subtracted a mask of pixels that likely contain a volume fraction of GM. This mask is obtained by thresholding the corresponding partial volume map at 0.05, and it ensures components are not extracted from voxels containing a minimal fraction of GM. Finally, these masks are resampled into BOLD space and binarized by thresholding at 0.99 (as in the original implementation). Components are also calculated separately within the WM and CSF masks. For each CompCor decomposition, the k components with the largest singular values are retained, such that the retained components’ time series are sufficient to explain 50 percent of variance across the nuisance mask (CSF, WM, combined, or temporal). The remaining components are dropped from consideration. The head-motion estimates calculated in the correction step were also placed within the corresponding confounds file. The confound time series derived from head motion estimates and global signals were expanded with the inclusion of temporal derivatives and quadratic terms for each^87^. Frames that exceeded a threshold of 0.5 mm FD or 1.5 standardised DVARS were annotated as motion outliers. All resamplings can be performed with a *single interpolation step* by composing all the pertinent transformations (i.e. head-motion transform matrices, susceptibility distortion correction when available, and co-registrations to anatomical and output spaces). Gridded (volumetric) resamplings were performed using antsApplyTransforms (ANTs), configured with Lanczos interpolation to minimize the smoothing effects of other kernels^88^. Non-gridded (surface) resamplings were performed using mri_vol2surf (FreeSurfer).

Many internal operations of *fMRIPrep* use *Nilearn* 0.8.1(RRID:SCR_001362)^89^, mostly within the functional processing workflow. For more details of the pipeline, see the section corresponding to workflows in *fMRIPrep*’s documentation.

#### Regions of interest

We used the Harvard-Oxford subcortical atlas (distributed with the *Nilearn* Python library, thresholded at 50% probability) to define the hippocampus region of interest (ROI), following past studies of hippocampal cognitive maps ^90,91^. We further defined anterior and posterior divisions of hippocampus based on the widely-used MNI coordinate, following Poppenk et al.^92^. The entorhinal cortex ROI was defined using the Juelich atlas (distributed with the *Nilearn* Python library, thresholded at 50% probability), following past studies of entorhinal cognitive maps ^27^. All ROI masks were then lateralized using MNI coordinate, into the corresponding left and right regions to produce six ROIs that were reverse-normalized from MNI space to participants’ native T1w spaces using *ANTS* ^93^, based on a set of transformations computed by *fMRIPrep* during preprocessing.

#### fMRI data analysis

We estimated first-level generalized linear models (GLMs) using the Nilearn library in Python (RRID:SCR_001362)^94^. As our main analyses are cross-validated across runs, we estimated a GLM per run, per participant. To mimic the default settings of the SPM Matlab toolbox, all GLMs used the ‘SPM + time derivative’ HRF model, grand-mean signal scaling, and the AR-1 noise model. Given our interest in information contained in fine-grained spatial patterns of neural activity, all GLMs were fit to images in native T1w spaces, and were left unsmoothed. All GLMs included nuisance regressors for the standard six head-motion parameters (i.e., three rigid-body translations and three rotations), their derivatives, their squares, and their squared derivatives. We also included additional nuisance regressors for the global signal, the first six anatomical CompCor components, and slow-drift cosine regressors. All nuisance variables were computed by fMRIPrep during preprocessing. From each GLM, we computed contrasts for every unique network member displayed to participants in the scanner, resulting in 22 whole-brain beta maps and one map of residuals mean square error per run.

Using the outputs from these first-level GLMs, we performed Representational Similarity Analysis (RSA)^95^ with the Python implementation of RSA Toolbox ^96^. To obtain unbiased and reliable estimates of neural similarity, we computed cross-validated Mahalanobis (i.e., ‘crossnobis’) distances ^50^. Given that cross-validating estimates of neural pattern similarity across runs produces an unbiased measure, the crossnobis estimator tends to be more reliable than other estimators through its use of multivariate noise normalization, which leverages the first-level GLM residuals to account for noise covariance. The ‘raw’ crossnobis estimates reflect dissimilarity, so in our statistical analyses, we applied a sign-flip such that more positive (less negative) values now reflect greater similarity for convenience.

### Computational modelling

#### Models of representation

We considered four candidate models of representation: Perfect Observation, Limited Observation, Katz Communicability (Katz), and Successor Representation (SR). The common assumption across these models is that people can acquire information about others’ relationships in the network. Using participants’ responses in the friendship survey, we constructed a matrix, *M*, of size N × N, where N corresponds to the number of network members (N = 187) and the value in a cell of this matrix, *M*(*i*, *j*), indicates whether members *i* nominated member *j* as their friend (0: not friend; 1:friend). Because not all friendship nominations are reciprocated, we only use mutually identified friendships to construct the ground-truth structure of the social network, represented by the adjacency matrix, *A*. The Perfect Observation model then simply assumes that participants’ mental representations of the network reflect this ground-truth structure, *A*.

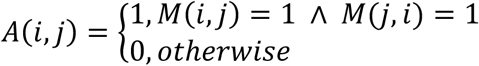

In contrast, the Limited Observation model instead assumes that people are constrained in their ability to acquire information about all the mutual friendships in the network. Here, we note two constraints. First, we expect that people are most likely to observe social interactions occurring within their immediate ‘social circle’ and are less and less likely to observe more distal relations ^45^, and estimate a parameter, *ω* ∈ [0,1], to quantify how strongly observations are affected by network distance, *d*. When *ω* → 0, participants only have information about their own friendships, whereas as *ω* → 1, they possess greater knowledge about relationship between dyads increasingly further away from them. Second, we consider the possibility that these observations do not depend on the strict reciprocity of relations, both between the observer and the dyad observed, and between members of the dyad themselves. Specifically, we consider that a person—e.g., Frank—is likely to observe someone they identify as their friend—e.g. Jack, to gain insight into who Jack identifies as their friend, even if Jack does not reciprocate Frank’s friendship. Through this, Frank possibly finds out that Jack is friends with Max, and to a lesser extent, also who Max identifies as their friend.

Together, this produces a systematically biased representation, *A’*, that is largely dependent on the outbound edges of the directed social network, *M*, where observations of the relationship between network members *i* and *j*, depends first and foremost on if either of them identifies the other as a friend. If neither do, the model assumes that people have no evidence of a relationship between the dyad. If only one member of the dyad identifies the other as their friend, the likelihood of the observation depends on the number of outbound edges between the participant and the member who nominated the other. If both members of the dyad mutually identify the other as their friend, the likelihood of the observation depends on the number of outbound edges that separate the participant from the closer member of the dyad, Additional model comparisons suggest that this assumption that people’s observations of the network depend on outbound edges better captures participants’ Network Knowledge than models based on mutual edges and inbound edges (see Supplementary Note S1).

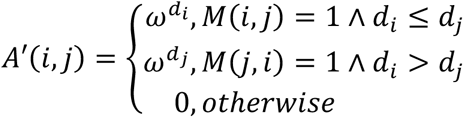

Our models of abstraction—i.e., Katz and SR—similarly assume that people are limited in their ability to directly acquire information about all the mutual friendships like the Limited Observation model, but additionally abstract over these observations to generate inferences based on the multi-step connections between each pair of network members. The Katz model abstracts directly over the biased adjacency matrix, *A’*, to produce the Katz communicability matrix, KC^35,38–41^.

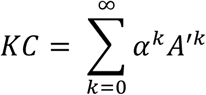

In plain terms, when given the ground-truth adjacency matrix, *A*, we can compute the powers of this matrix *A*^k^ which produces a matrix that indexes the number of k-step relations separating each pair of network members (or alternatively, the number of k-step paths connecting the pair). When applied to the biased adjacency matrix, *A’*, *A’*^k^ analogously reflects the extent of k-step connectivity between two pairs, qualified by the participants’ limited ability to observe each step. The Katz communicability is thus the weighted sum of all the multi-step connections between each pair of network members, where each multi-step connection is discounted by α based on the number of steps it comprises. Because α has a theoretical upper bound equal to the inverse of the maximum eigenvalue of the matrix, *A’*, we can re-express the Katz communicability as:

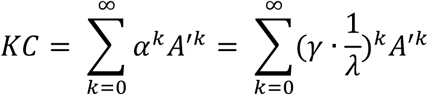

where λ is the maximum eigenvalue of *A’* and γ ∈ [0,1) such that γ indexes the degree to which a participant integrates (i.e., abstracts over) increasing number of steps when representing pairwise associations in the network. When γ → 0, the representation only contains direct observations of pairwise relations consistent with memorization. As γ → 1, the representation becomes increasingly abstract, reflecting longer-range connections between a pair of individuals.

The SR, much like the Katz, abstracts over the observable pairwise relations in the network^20,30,43,44^. However, instead of abstracting directly over the biased adjacency matrix, *A’*, it first normalizes it into a transition matrix, *T’*, which represents the relative probability that member *i* is friends with member *j*, as opposed to all other network members in a network of size *N*.

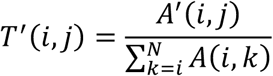

The SR is then computed as the weighted sum of the multi-step transition probabilities between each pair of network members, where each multi-step transition is discounted by γ ∈ [0,1) based on the number of steps it comprises.

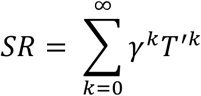

Correspondingly, because both the Katz and SR take the form of a geometric sum of an infinite series of square matrices, we can use the closed-form solution to obtain these representations given some value of γ and the biased adjacency/transition matrix^20^:

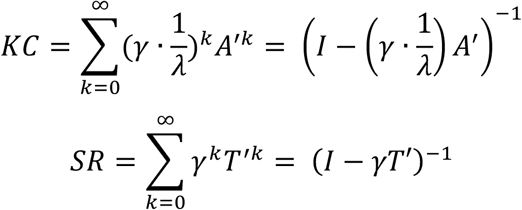

where *I* represents the identity matrix and the superscript (−1) represents the inversion of the matrix.

#### Model-fitting

We estimated participants’ subjective representations of the friendship network, *R*, using these four models by combining each of these representational formats with a logistic choice function to predict participant’s judgments about whether network members *i*, and *j* are friends in the Network Knowledge Task.

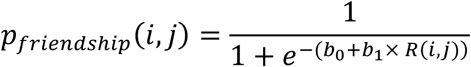

This allowed us to compute the likelihood of participants’ responses on all the trials in the Network Knowledge Task given some set of parameter values for each of the representational formats. Consequently, the Perfect Observation model has two free parameters (b_0_, b_1_), the Limited Observation model has three (b_0_, b_1_, ω), while Katz and SR each have four (b_0_, b_1_, ω, γ).

We estimated these parameters hierarchically using Differential Evolution Monte-Carlo Markov Chains (DEMCMC)^97,98^. In short, this approach uses the likelihood function of different computational models given the data to iteratively construct the posterior distribution of model parameters for each participant by proposing new sets of model parameter values and evaluating them using the Metropolis-Hastings algorithm. It then simultaneously constructs the posterior distribution of population hyperparameters given the current estimates of all participants in the same way through iteratively proposing and evaluating potential sets of hyperparameters.

For each model with k free parameters, we ran 3× k chains in parallel for 50,000 samples after a burn in of 100,000 (for all models except SR which required 150,000), thinning the samples by a factor of five. This yielded a total of k × 30,000 samples to construct the posterior distribution over parameter estimates. We additionally implemented a probabilistic migration step, *α* = 0.005, with every MCMC step in place of the differential evolution to improve chain-mixing and convergence towards the high probability density region of the posterior distribution of parameters. The migration step cycles the positions of a subset of chains (N_migrate_ uniformly sampled from the total number of chains) such that the positions of chains {i,i+1,…j-1, j} were compared against {i+1, i+2,…,j, i} and evaluated based on the Metropolis-Hastings algorithm^97,99^.

We assumed that participant-level parameters were distributed with respect to a set of population hyperparameters. We set the priors for hyperparameters of the logistic choice function such that the means of the choice parameters b_0_ and b_1_ were normally distributed, N(0,5), and standard deviations were drawn from a half-normal distribution, HN(0,1). Participant-specific b_0_ and b_1_ were drawn from a normal distribution based on these hyperparameters. To account for the fact that values of ω are theoretically bounded between 0 and 1, we defined the priors for the population mean to be drawn from a truncated normal distribution with mean = 1 and standard deviation = .5, TN(1,.5). This instantiates a conservative null hypothesis that people have perfect observation of far-away friendships—in contrast to our hypothesis that people have a limited ability to observe these friendships. Priors for the population standard deviation of ω was defined as a half-normal distribution, HN(0,.5). Participant-specific ω was then assumed to be drawn from a truncated-normal distribution bounded between 0 and 1 with mean and standard deviations corresponding to the population hyperparameters. Similarly, to account for the fact that values of γ are also bounded between 0 and 1, we likewise defined the priors for the population mean to be drawn from a truncated normal distribution with mean 0 and standard deviation .5, TN(0,.5). This in turn instantiates a conservative null hypothesis that people do not engage in any abstraction—in contrast to our hypothesis that people rely on multi-step abstraction. Priors for the population standard deviation of γ was defined as a half-normal distribution, HN(0,.5). Participant-specific γ was also then assumed to be drawn from a truncated-normal distribution bounded between 0 and 1 with mean and standard deviations corresponding to the population hyperparameters for γ.

Models were assessed to have converged to the posterior distribution as assessed by the Gelman-Rubin statistic for all hyperparameters and participant-specific parameters: *max*(R-hat) < 1.01^100,101^. Best-fit values of model parameters were extracted from the posterior distributions based on the *mean* estimate. Candidate models were recoverable based on simulated data generated from these estimates (see Supplementary Fig S1-S4). Using these estimates, we then reconstruct participant’s subjective representations of social ties within the network as a predictor of neural pattern similarity in subsequent analyses, and to simulate the responses they would make on the Friendship Knowledge Task as posterior predictive checks.

#### Posterior predictive checks

To assess the overall fit of each computational model, we first computed the analytic probability that a participant would identify a given pair of network members as friends in the Network Knowledge Task for every participant and each pair of network members based on the best-fitting parameters from each model. Next, we binned participants’ responses across trials based on their familiarity with the dyad (minimum graph distance), and the relationship between each member of the dyad, and averaged the simulated probability and observed proportion of participants’ judgments that the dyad are friends. We then calculated the within-participant Pearson correlation between the simulated and observed responses across these different levels of the participants’ familiarity with the dyad and the dyad’s relationship and conducted a one-sample t-test of these correlation coefficients to determine if the simulated responses quantitatively captured the observed patterns of participants’ behavior.

#### Model comparison

To formally compare our different models of representation, we first computed the Watanabe-Akaike Information Criterion (WAIC)^102–104^ of each model by randomly sampling 1,000 sets of model parameters from the estimated posterior distribution of each participant. We then evaluated the likelihood of each response, *y*, the participant made given each set of model parameters, *θ*, averaged across all *S* number of simulations and log-transformed them to obtain the log predictive probability of a single trial, *i*. Summing across all *n* number of trials then produces the log pointwise predictive density, which captures the overall fit of the model to the data.

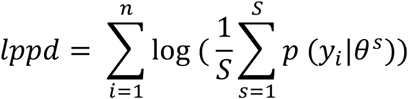

To account for potential overfitting, the WAIC imposes a penalty on the models that can be approximated by the variance in predictions generated by the parameter sets sampled from the posterior distribution summed across observed responses.

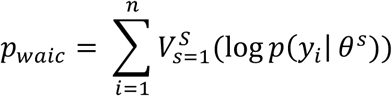

The resulting value is then multiplied by a factor of –2 to adhere to the deviance scale consistent with the AIC.

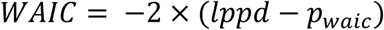

After computing the WAIC for each participant and each model, we then derived three indices of relative model fit for comparison. First, we transformed WAIC for each model into Watanabe-Akaike weights for each participant to represent the relative evidence for each model in capturing the participant’s data^105^. Second, we calculated the protected exceedance probability which indexes the probability that each model best accounts for participants’ behavior across the entire sample^106^. Finally, we also obtained the overall group level WAIC by summing the individual-level WAICs across all participants to compute the Watanabe-Akaike weight for each model at the group-level.

### Software

All computational modelling was conducted in Python v3.9.18. using *<u>numpy</u>* v1.26.4^107^, *pandas* v2.2.1^108^, *numba* v0.59.0^109^, *scipy* v1.12.0^110^, *arviz* v0.16.0^111^, and *spyder* v5.5.1 in *Anaconda* v2-2.4.0^112^. All statistical analyses were conducted in RStudio v2023.12.1+402 (RStudio Team, 2015) with R v4.3.3 (R Core Team, 2017). We fitted linear mixed effects models to continuous data with degrees of freedom estimated using the Satterthwaite method, and logistic mixed-effects model to binary data using the *lme4* package v1.1-35.2^113,114^, with maximal random effects where possible. Ordered beta regressions^61^ were fitted to continuous data bounded between 0 and 1 using *glmmTMB* v1.1.10^115^. Simple slopes were extracted using the *marginaleffects* package v0.21.0^116^. Plots of participant data and model predictions were generated using the *ggplot2* v3.5.0^117^and *ggeffects* v1.5.1^118^ packages. Social network analyses were also conducted in Rstudio with R using the *igraph* v2.0.3^119,120^ and *tidygraph* v1.3.1^121^ packages. Visualization of the network was plotted using *ggraph* v2.2.1^122^. Software for fMRI processing is reported above in the section on functional neuroimaging. Stimuli presentation and the recording of participants’ responses were supported by *Qualtrics* in the Friendship Surveys and Network Knowledge Task, *JsPsych* v7.1^123^ in the Information Flow Task, and *PsychToolBox*-3^124^ in *MATLAB* (R2017b)^125^ for the functional neuroimaging session.

## Code & Data Availability

All necessary data and code necessary to reproduce the results are openly accessible at https://osf.io/adm8v/?view_only=8e939a2154444673a26d6a3419bf5a0a. All data provided is de-identified and labeled with randomly assigned pseudonyms to facilitate matching across datasets.

## Acknowledgements

We would like to thank Isabella Aslarus, Yi-Fei Jerry Hu, Elizabeth Duchan, Maya Mazumder, Kayleigh Danowski, Jonathan Palfy, Vera Poyraz, Samantha Shulman, Sofia Vaca Narvaja, and Jenny Wang for their indispensable assistance in data collection. Part of this research was conducted using computational resources and services at the Center for Computation and Visualization, Brown University. Advanced access to these computing resources was supported by NIH award 1S10OD025181. This work is supported by the National Science Foundation award 2123469 to O.F.H. and A.B.

## Author contributions

Y.Y.T. and J.Y.S. contributed equally to this work. A.B. and O.F.H. contributed equally to this work. Conceptualization: Y.Y.T., J.Y.S., A.B. and O.F.H. Formal Analysis: Y.Y.T. and J.Y.S. Funding Acquisition: A.B. and O.F.H. Investigation: J.Y.S. and A.X. Methodology: Y.Y.T., J.Y.S., A.X., A.B. and O.F.H. Software Y.Y.T., J.Y.S. and A.X. Supervision: A.B. and O.F.H. Validation: Y.Y.T. and A.X. Writing – original draft: Y.Y.T., J.Y.S., A.B. and O.F.H. Writing – review and editing: Y.Y.T., J.Y.S., A.X., A.B. and O.F.H.

## Supplementary Information

**Supplementary Table S1:** Behavioral analysis of Network Knowledge Task.

| **Predictor** | **Estimate** | **95% CI** | **Z** | **p** |
| --- | --- | --- | --- | --- |
| Intercept | -3.841 | [-4.177, -3.505] | -22.436 | < .001 |
| Unfamiliarity | -0.359 | [-0.552, -0.166] | -3.652 | < .001 |
| Pairwise Friendship | 2.067 | [1.840, 2.295] | 17.823 | < .001 |
| Multi-step Relation | -0.882 | [-1.053, -0.712] | -10.145 | < .001 |
| Unfamiliarity × Pairwise Friendship | -1.011 | [-1.185, -0.837] | -11.373 | < .001 |
| Unfamiliarity × Multi-step Relation | 0.417 | [0.246, 0.588] | 4.786 | < .001 |

**Supplementary Table S2:** Associations between computational model parameters and Network Knowledge Accuracy.

| Predictor | Estimate | 95% CI | t | df | p |
| --- | --- | --- | --- | --- | --- |
| Intercept | 1.094 | [0.986, 1.203] | 19.995 | 95 | < .001 |
| $\omega$ | 0.549 | [0.072, 1.025] | 2.286 | 95 | 0.024 |
| $\gamma$ | 1.162 | [0.160, 2.163] | 2.303 | 95 | 0.023 |
| $\gamma^2$ | -8.723 | [-14.957, -2.489] | -2.778 | 95 | 0.007 |
| Response bias | -0.122 | [-0.182, -0.062] | -4.056 | 95 | < .001 |

Note: A simple linear model was fitted to participants' accuracy in judging true friendships, measured by d-prime, as a function of the (mean-centered) best-fitting model parameters from the Katz model. The parameter $\omega$ captures the participant's ability to acquire direct information about pairwise friendships more distal to them. The parameter $\gamma$ captures the degree to which participants' abstract over these observations. The response bias is the intercept of the logistic choice function in the computational model that indicates their overall response tendency to say that a pair was friends. Model comparison with a nested model that drops the quadratic effect of $\gamma$ found that the nested model fit the data worse ( $\Delta\text{AIC} = 5.810 > 2$ )<sup>126</sup>, suggesting that the quadratic effect of $\gamma$ explained a significant amount of variance in why some participants were more accurate than others. The parameter value corresponding to the vertex of the parabola produced by the quadratic effect was computed based on the algebraic formula $-b/2a$ , where $a$ is the quadratic estimate and $b$ is the linear estimate. Because the analysis was conducted on mean-centered parameter values, the real vertex ( $\gamma = 0.682$ ) was obtained by adding the sample's mean value of $\gamma$ to the evaluation of the algebraic expression.

**Supplementary Table S3:**
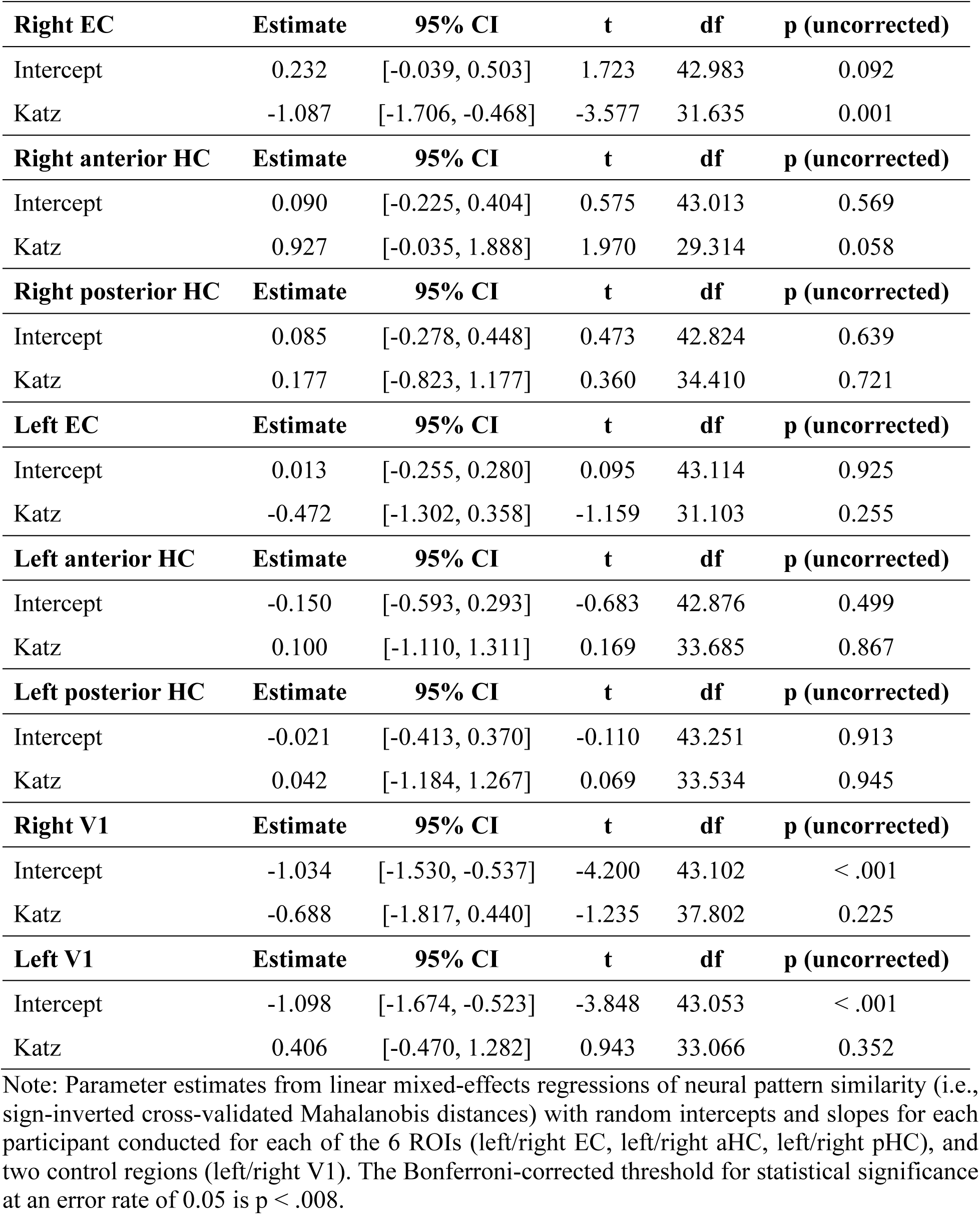
Neural encoding of Katz Communicability.

**Supplementary Table S4:**
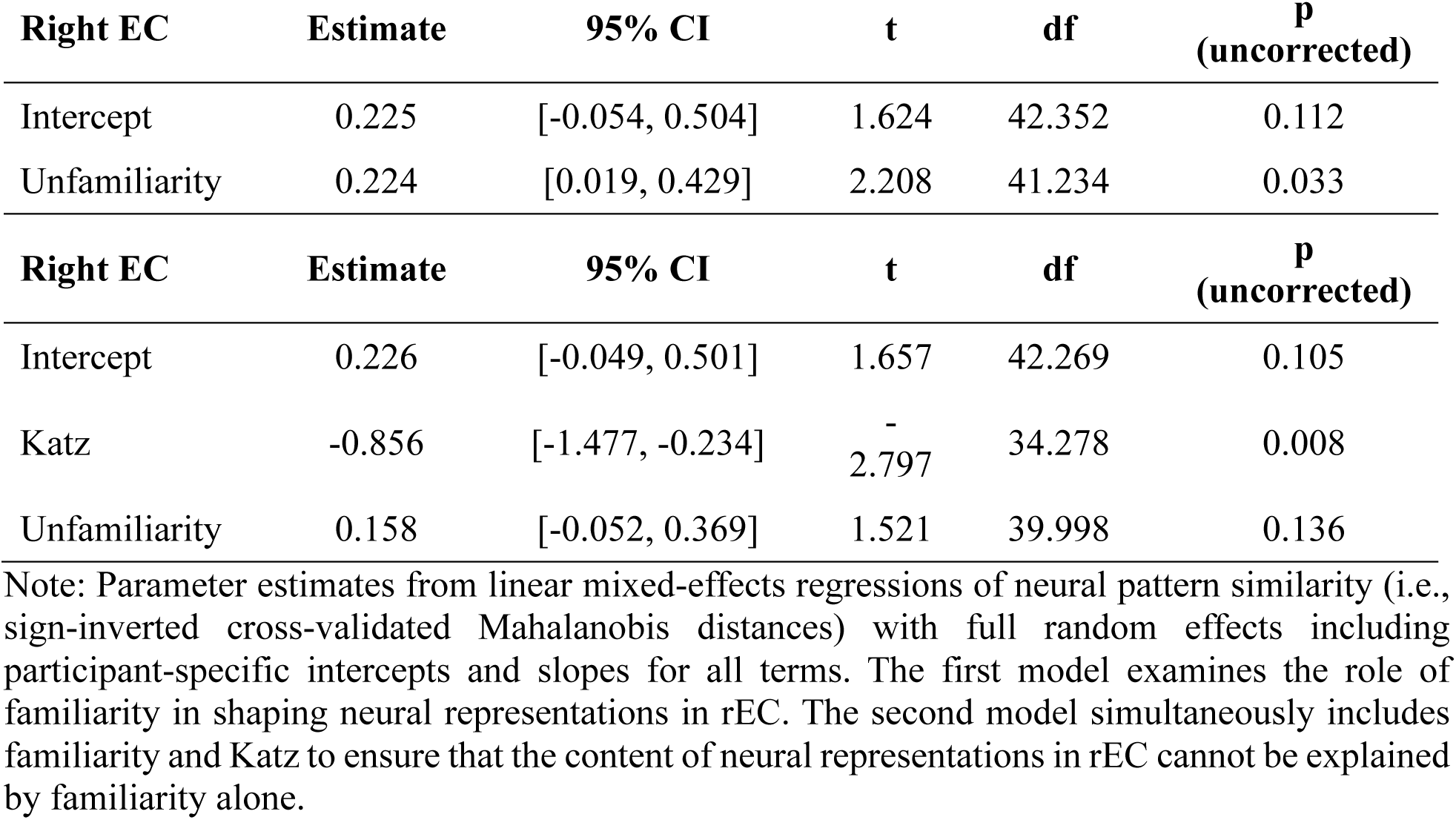
Neural encoding of Katz Communicability controlling for familiarity in rEC.

**Supplementary Table S5:** Neural encoding of Katz Communicability controlling for Observations in rEC and raHC.

| **Right EC** | **Estimate** | **95% CI** | **t** | **df** | **p (uncorrected)** |
| --- | --- | --- | --- | --- | --- |
| Intercept | 0.229 | [-0.044, 0.501] | 1.691 | 42.919 | 0.098 |
| Katz | -1.495 | [-2.408, -0.581] | -3.312 | 38.320 | 0.002 |
| Observations | 0.056 | [-0.033, 0.145] | 1.239 | 144.476 | 0.217 |

| **Right anterior HC** | **Estimate** | **95% CI** | **t** | **df** | **p (uncorrected)** |
| --- | --- | --- | --- | --- | --- |
| Intercept | 0.072 | [-0.253, 0.398] | 0.449 | 41.909 | 0.656 |
| Katz | 0.924 | [-0.958, 2.807] | 1.039 | 16.390 | 0.314 |
| Observations | -0.023 | [-0.195, 0.149] | -0.292 | 13.124 | 0.774 |

Note: Parameter estimates from linear mixed-effects regressions of neural pattern similarity (i.e., sign-inverted cross-validated Mahalanobis distances) with full random effects including participant-specific intercepts and slopes for all terms, except the random-slope for observations from the model of rEC patterns was dropped due to non-convergence and the estimated between participant variance approximating zero.

**Supplementary Table S6:** Parsing the content of rEC neural codes using residualized predictors of Katz and Observations.

| **Right EC** | **Estimate** | **95% CI** | **t** | **df** | **p**<br>**(uncorrected)** |
| --- | --- | --- | --- | --- | --- |
| Intercept | 0.228 | [-0.045, 0.501] | 1.685 | 42.986 | 0.099 |
| Residualized Inferences | -2.062 | [-3.357, -0.767] | -3.268 | 26.867 | 0.003 |
| Observations | -0.096 | [-0.158, -0.034] | -3.180 | 30.293 | 0.003 |
| Intercept | 0.232 | [-0.040, 0.504] | 1.719 | 43.066 | 0.093 |
| Inferences | -1.096 | [-1.710, -0.482] | -3.643 | 31.136 | < .001 |
| Residualized Observations | 0.173 | [0.014, 0.332] | 2.197 | 37.672 | 0.034 |

Note: Parameter estimates from linear mixed-effects regressions of neural pattern similarity (i.e., sign-inverted cross-validated Mahalanobis distances) with full random effects including participant-specific intercepts and slopes for all terms. All random effects were specified to be independent as inclusion of the random-effects covariances with the residualized variables led to non-convergence of the models. Because Katz integrates both the directly observed one-step relations and multi-step relations, we can decompose the metric into direct observations of the one-step connection: $A'$ , and the inferred multi-step connectivity: $Katz - \gamma * A' / \lambda$ . However, these observations and inferences remain moderately correlated. Thus, to ascertain which explains the greatest proportion of variability in neural patterns, we run two models residualizing one metric against the other and vice versa. In the first model, we residualize Inferences against Observations, effectively attributing all shared variance between them to the Observations variable. In the second model, we residualize Observations against Inferences, attributing all shared variance to the Inferences variable. Consistent effects of inferences across the models suggest that the rEC robustly encodes the inferential component of Katz.

**Supplementary Table S7:** Parsing the content of raHC neural codes using residualized predictors of Katz and Observations.

| **Right anterior HC** | **Estimate** | **95% CI** | **t** | **df** | **p (uncorrected)** |
| --- | --- | --- | --- | --- | --- |
| Intercept | 0.090 | [-0.223, 0.404] | 0.581 | 43.009 | 0.564 |
| Residualized Inferences | 0.947 | [-1.980, 3.874] | 0.676 | 19.505 | 0.507 |
| Observations | 0.077 | [-0.010, 0.164] | 1.825 | 25.614 | 0.080 |
| **Right anterior HC** | **Estimate** | **95% CI** | **t** | **df** | **p (uncorrected)** |
| Intercept | 0.090 | [-0.225, 0.405] | 0.577 | 43.033 | 0.567 |
| Inferences | 0.931 | [-0.041, 1.903] | 1.955 | 30.324 | 0.060 |
| Residualized Observations | -0.004 | [-0.315, 0.306] | -0.029 | 28.291 | 0.977 |

**Supplementary Table S8:**
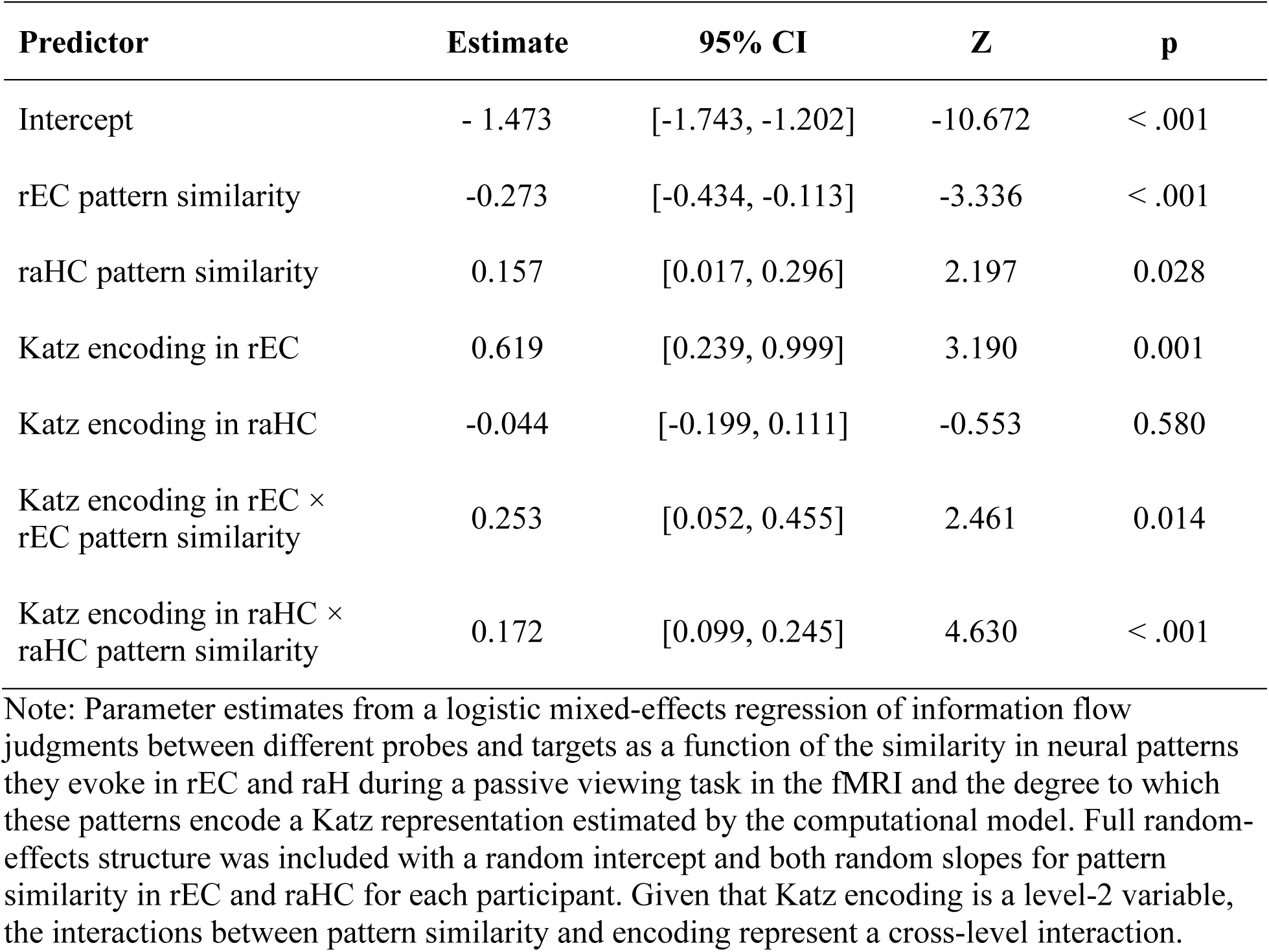
rEC and raHC encoding of relational maps support information flow judgments.

**Supplementary Table S9:**
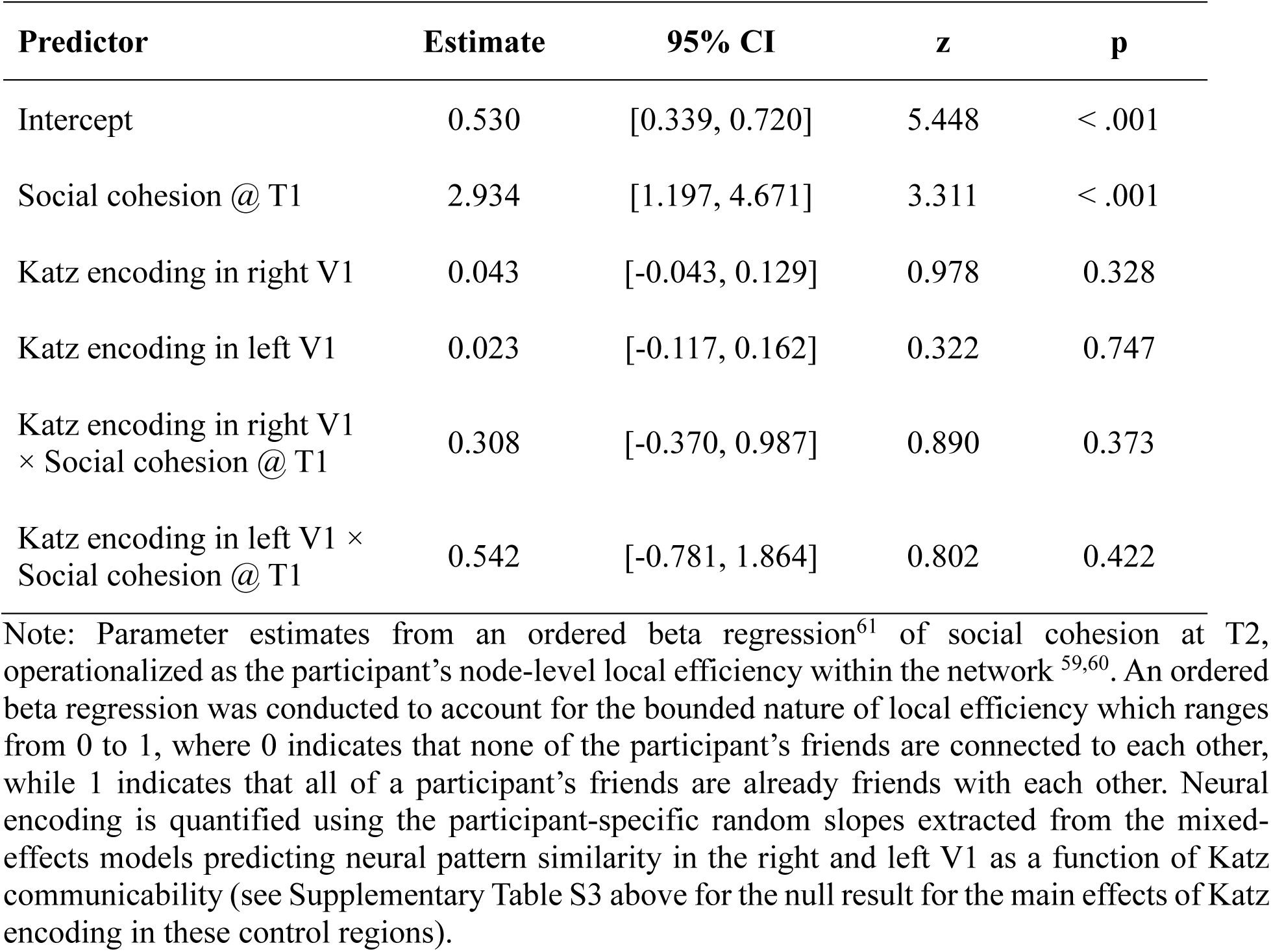
Control analysis examining whether strategic social brokerage can be explained by Katz encoding in V1.

| Predictor | Estimate | 95% CI | z | p |
| --- | --- | --- | --- | --- |
| Intercept | 0.530 | [0.339, 0.720] | 5.448 | < .001 |
| Social cohesion @ T1 | 2.934 | [1.197, 4.671] | 3.311 | < .001 |
| Katz encoding in right V1 | 0.043 | [-0.043, 0.129] | 0.978 | 0.328 |
| Katz encoding in left V1 | 0.023 | [-0.117, 0.162] | 0.322 | 0.747 |
| Katz encoding in right V1<br>× Social cohesion @ T1 | 0.308 | [-0.370, 0.987] | 0.890 | 0.373 |
| Katz encoding in left V1 ×<br>Social cohesion @ T1 | 0.542 | [-0.781, 1.864] | 0.802 | 0.422 |

Note: Parameter estimates from an ordered beta regression<sup>61</sup> of social cohesion at T2, operationalized as the participant's node-level local efficiency within the network<sup>59,60</sup>. An ordered beta regression was conducted to account for the bounded nature of local efficiency which ranges from 0 to 1, where 0 indicates that none of the participant's friends are connected to each other, while 1 indicates that all of a participant's friends are already friends with each other. Neural encoding is quantified using the participant-specific random slopes extracted from the mixed-effects models predicting neural pattern similarity in the right and left V1 as a function of Katz communicability (see Supplementary Table S3 above for the null result for the main effects of Katz encoding in these control regions).

**Supplementary Table S10:** Strategic social brokerage referencing changes in cohesion to immediately after neuroimaging rather than prior to neuroimaging.

| Predictor | Estimate | 95% CI | Z | p |
| --- | --- | --- | --- | --- |
| Intercept | 0.582 | [0.427, 0.738] | 7.341 | < .001 |
| Social cohesion @ T1.5 | 2.296 | [1.408, 3.185] | 5.064 | < .001 |
| Katz encoding in rEC | -0.384 | [-0.587, -0.181] | -3.714 | < .001 |
| Katz encoding in raHC | 0.129 | [0.024, 0.234] | 2.418 | 0.016 |
| Katz encoding in rEC ×<br>Social cohesion @ T1.5 | 1.643 | [0.202, 3.084] | 2.234 | 0.025 |
| Katz encoding in raHC ×<br>Social cohesion @ T1.5 | -0.250 | [-0.939, 0.439] | -0.712 | 0.477 |

Note: Parameter estimates from a simple ordered beta regression of social cohesion at T2 similar to in Table 1 in the main text, except that we alternatively control for social cohesion of the participants' friends using a measure of the network taken within a two-week period right after the neuroimaging session rather than prior (T1.5; N = 42, one neural participant did not return for this survey right after). This analysis ensures that the association between neural encoding and increases in social cohesion cannot simply be explained by changes in friendships between our survey at T1 and the neuroimaging session, that is already reflected in participants' neural representations. Here, we find more robust effects of rEC and raHC encoding in predicting increases in social cohesion, although for the rEC, this is qualified by an interaction with levels of cohesion at T1.5. This interaction suggests that encoding strength predicted increases in social cohesion for participants' whose communities were not very cohesive, but not for participants whose community was already highly cohesive. Further examination reveals that this is likely due to potential ceiling effects, since the maximum local efficiency of participants at T1 is 0.836, whereas maximum local efficiency of participants at T1.5 is the theoretical maximum of 1.

**Supplementary Table S11:** Strategic social brokerage controlling for changes in immediate community.

| Predictor | Estimate | 95% CI | Z | p |
| --- | --- | --- | --- | --- |
| Intercept | 0.650 | [0.462, 0.839] | 6.755 | < .001 |
| Social cohesion @ T1 | 2.988 | [1.474, 4.502] | 3.868 | < .001 |
| Katz encoding in rEC | -0.267 | [-0.492, -0.043] | -2.336 | 0.019 |
| Katz encoding in raHC | 0.111 | [0.009, 0.212] | 2.139 | 0.032 |
| Similarity in friendships from T1 to T2 | -0.421 | [-1.412, 0.571] | -0.831 | 0.406 |
| Katz encoding in rEC × Social cohesion @ T1 | -0.116 | [-2.477, 2.245] | -0.096 | 0.923 |
| Katz encoding in raHC × Social cohesion @ T1 | -0.318 | [-1.442, 0.805] | -0.556 | 0.578 |
| Katz encoding in rEC × Similarity in friendships | -0.510 | [-1.959, 0.939] | -0.690 | 0.490 |
| Katz encoding in raHC × Similarity in friendships | -0.461 | [-1.093, 0.172] | -1.427 | 0.154 |

Note: Parameter estimates from as simple ordered beta regression of social cohesion at T2 similar to in Table 1 in the main text, except that we additionally control for the similarity in participants' friends across the two time points (T1 and T2). Similarity was quantified using Jaccard similarity which indicates the proportion of friends participants reported having being common across the two time points. Given that our effects were not modulated by similarity, our effects could not have been selectively driven by participants who left their communities to join other highly cohesive communities across the two time points (i.e., people with low friendship similarity across times).

**Supplementary Table S12:** Strategic social brokerage controlling for degree of abstraction.

| Predictor | Estimate | 95% CI | Z | p |
| --- | --- | --- | --- | --- |
| Intercept | 0.577 | [0.415, 0.738] | 7.009 | < .001 |
| Social cohesion @ T1 | 2.900 | [1.508, 4.291] | 4.085 | < .001 |
| Katz encoding in rEC | -0.242 | [-0.457, -0.028] | -2.216 | 0.027 |
| Katz encoding in raHC | 0.052 | [-0.049, 0.152] | 1.011 | 0.312 |
| Degree of abstraction: $\gamma$ | 0.357 | [-1.103, 1.818] | 0.480 | 0.631 |
| Katz encoding in rEC ×<br>Social cohesion @ T1 | -0.795 | [-3.042, 1.451] | -0.694 | 0.488 |
| Katz encoding in raHC ×<br>Social cohesion @ T1 | -0.575 | [-1.690, 0.539] | -1.012 | 0.312 |
| Katz encoding in rEC ×<br>Degree of abstraction | 0.038 | [-1.628, 1.703] | 0.045 | 0.964 |
| Katz encoding in raHC ×<br>Degree of abstraction | -1.259 | [-2.229, -0.288] | -2.542 | 0.011 |

Note: Parameter estimates from a simple ordered beta regression of social cohesion at T2 similar to in Table 1 in the main text, except that we additionally control for the abstractness of the maps encoded in rEC and raHC. This degree of abstraction is quantified using the parameter $\gamma$ estimated from our model. Given that participants in our sample overall engaged in at least moderate levels of abstraction (min $\gamma = 0.456$ , mean $\gamma = 0.605$ , max $\gamma = 0.830$ ), the robust main effect of Katz encoding in rEC emphasize that it is the spontaneous availability of these maps rather than the abstractness of these maps that predict social cohesion @ T2, suggesting that they may afford participants the ability to draw on the relational information encoded within to actively broker friendships within their community. In contrast, this analysis also reveals that encoding in the raHC predicts greater social cohesion @ T2 only when the maps encoded were relatively less abstract. We refrain from interpreting this given the lack of robustness of the raHC effects across model specifications.

**Supplementary Note S1: Alternative specifications of information acquisition about others’ friendships.**

Given that our study examined people’s cognitive and neural representations of their naturalistic social network, computational modelling of these processes required us to make assumptions about the information people could directly acquire about pairwise friendships between others. In our Perfect Observation model, we considered that our participants could obtain complete and veridical knowledge about all pairwise relationships. In contrast, our Limited Observation, and abstraction models built on these limited observations (i.e., Katz & SR) assume that people have a limited and biased access to information about the relationship between a pair of network members based on how distant they are to the pair ^31,45^. However, although we tend to assume that a fundamental feature of friendship is that it is reciprocal and thus define the network as that of mutual friendships, a non-trivial number of friendship nominations participants make in our study are in fact not reciprocated. Combined with our assumption that participants are limited and biased in their access to relational information, we consider the possibility that the relational information people acquire to construct their representations of the network, and how they acquire it depend on more than mutual friendships.

Here, we consider three possibilities through which people can access information about friendships based on their distance to the pair. First, access to relational information is based upon mutually identified friendship: People learn only about reciprocated friendships in the network through their reciprocated friendships. Alternatively, relational information is based on the identification of friendship by at least one member of the dyad: People acquire information about all the friendships others report and assume that they are reciprocal, even if they are not in fact. Thus, people’s representations could be built upon the directed network of friendship nominations rather than the reciprocal network. This in turn raises two possibilities: that information acquisition is limited by who members nominate (i.e., outbound edges), or that information acquisition is limited by who members are nominated by (i.e., inbound edges). To test these three distinct possibilities, we fitted three versions of all four of our candidate models that distinctly assume participants’ representations of the network mirror reciprocated, outbound or inbound friendship nominations.

Consistent with model comparison results reported in the main text, our Katz model based upon outbound nominations was the winning model among all possible candidates (average participant-level w(WAIC) = 0.309; pxp = .996; overall sample w(WAIC) > .999; Supplementary Fig S5A). Restricting this analysis to the three specific variants of the best-fitting Katz model to test the different assumptions of relational observation reveals that representations built upon outbound nominations were significantly better at capturing the data than representations built upon inbound and mutual nominations (average participant-level w(WAIC) = 0.506; pxp = .994; overall sample w(WAIC) > .999; Supplementary Fig S5B).

**Supplementary Fig S1.**
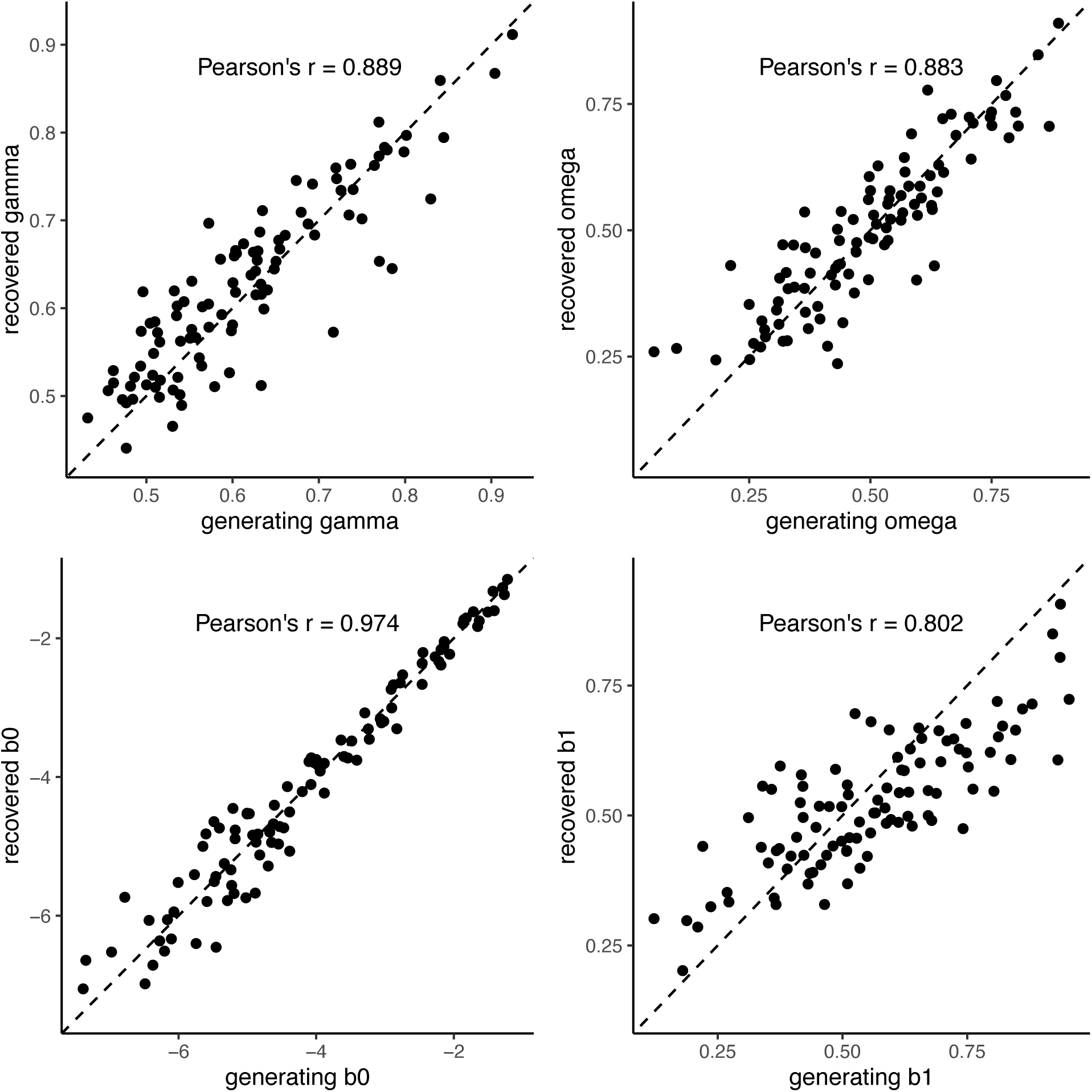
Model Recovery of the Katz model. Each point represents a single subject. The dashed diagonal line indicates perfect recovery. Model recovery was conducted by simulating responses for each participant based on best-fit parameters for this model and fitting the whole simulated dataset using the same model-fitting procedure to obtain participant parameters.

**Supplementary Fig S2.**
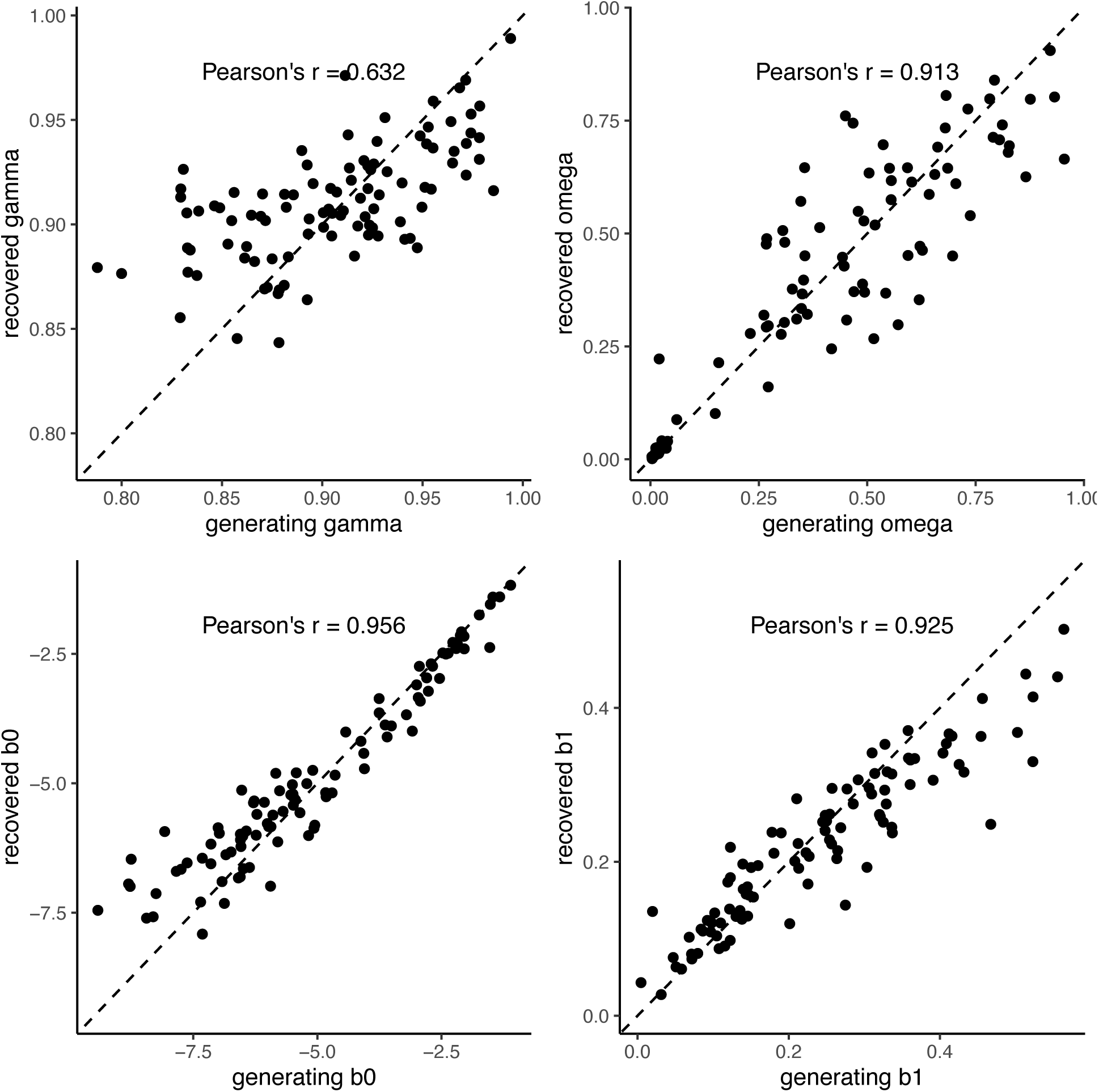
Model Recovery of the SR model. Each point represents a single subject. The dashed diagonal line indicates perfect recovery. Model recovery was conducted by simulating responses for each participant based on best-fit parameters for this model and fitting the whole simulated dataset using the same model-fitting procedure to obtain participant parameters.

**Supplementary Fig S3.**
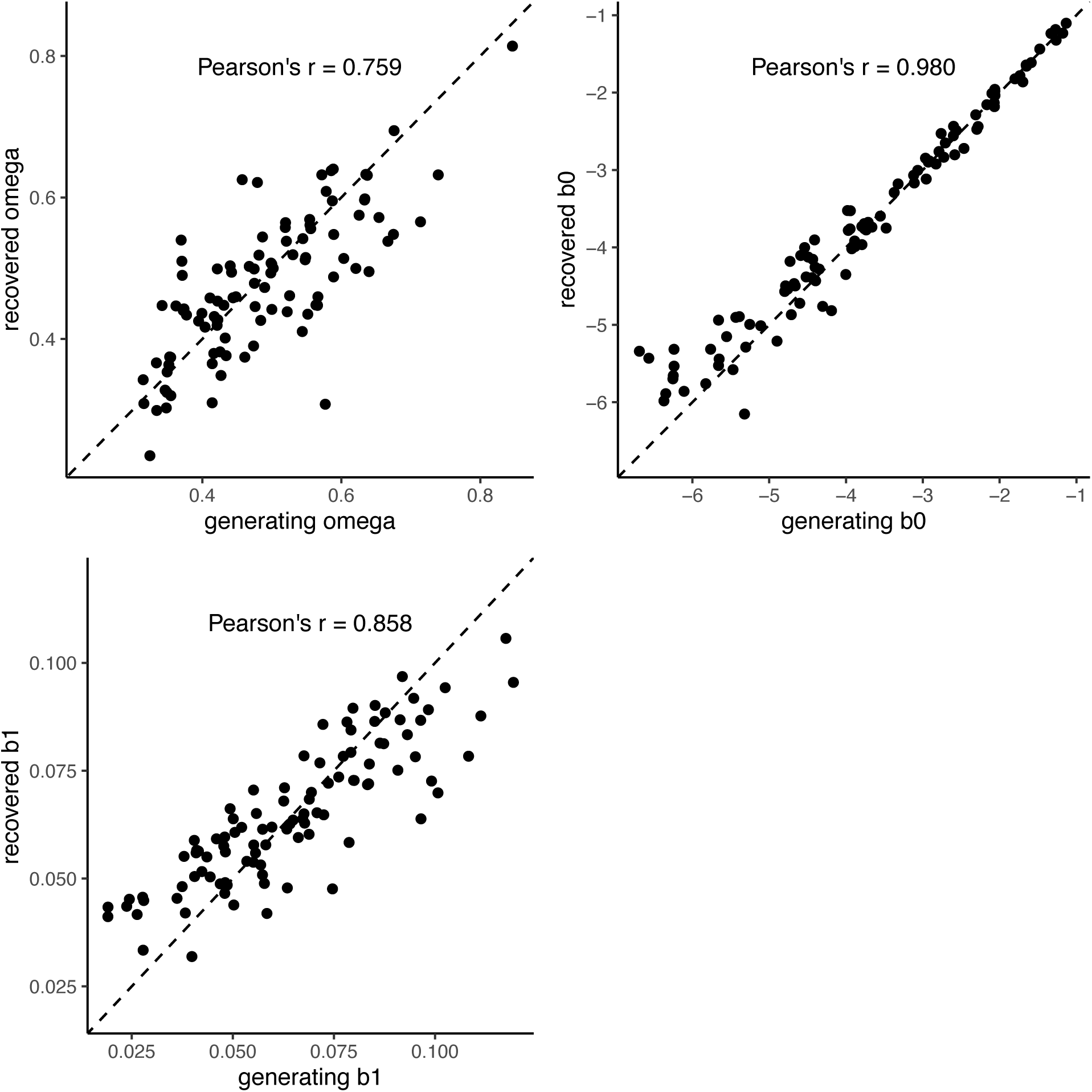
Model Recovery of the Limited Observation model. Each point represents a single subject. The dashed diagonal line indicates perfect recovery. Model recovery was conducted by simulating responses for each participant based on best-fit parameters for this model and fitting the whole simulated dataset using the same model-fitting procedure to obtain participant parameters.

**Supplementary Fig S4.**
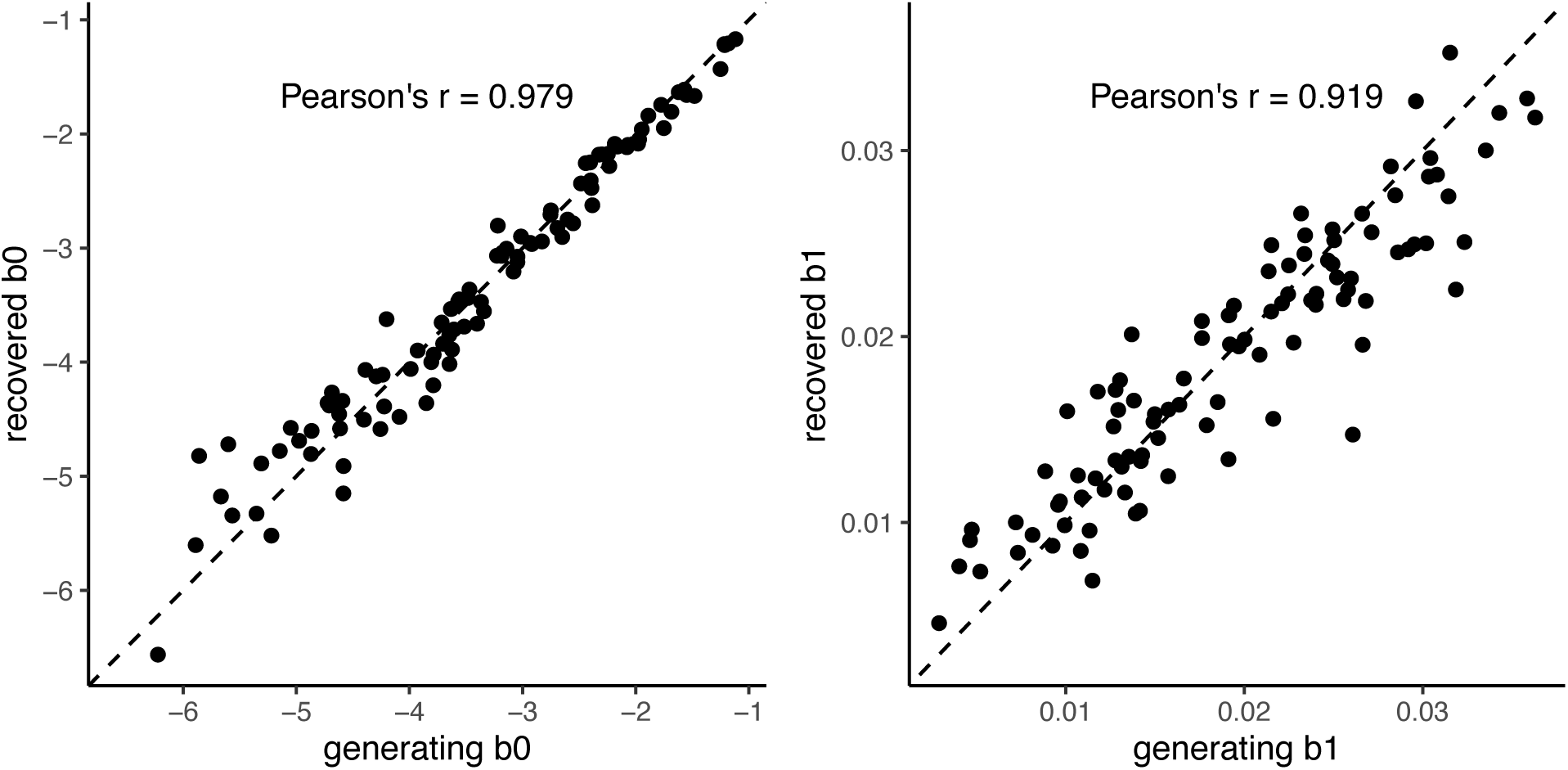
Model Recovery of the Perfect Observation model. Each point represents a single subject. The dashed diagonal line indicates perfect recovery. Model recovery was conducted by simulating responses for each participant based on best-fit parameters for this model and fitting the whole simulated dataset using the same model-fitting procedure to obtain participant parameters.

**Supplementary Fig S5.**
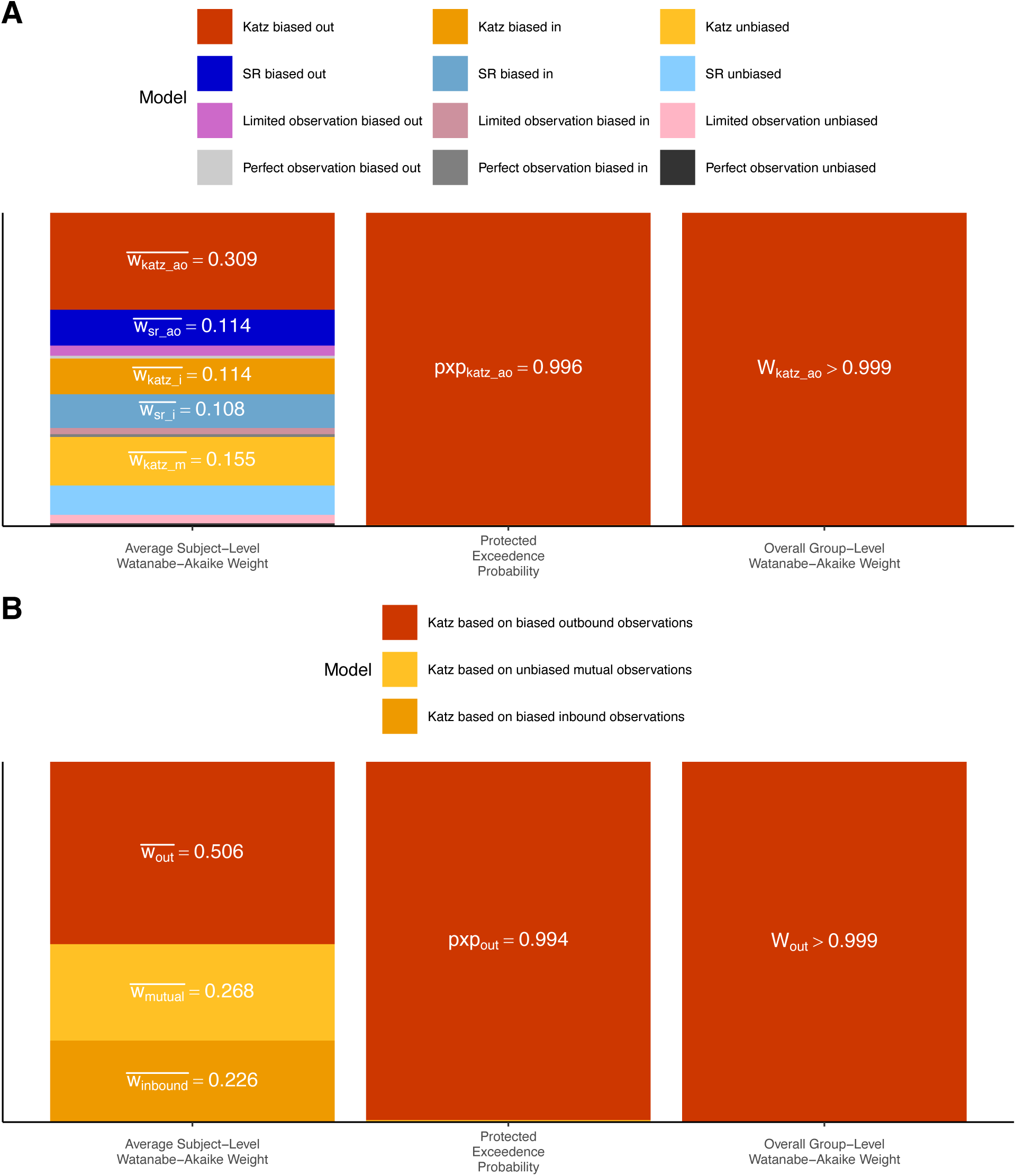
Model comparison. **A) All possible models of representation, each using three distinct assumptions of information dynamics within the network. B) Comparison of Katz models with distinct assumptions about relational observation.** Height of stacked bars indicate the relative evidence for each of the 12 candidate models in (A), and for each of the three variants of the best-fitting Katz model in (B) based on the average participant-level Watanabe-Akaike weight, protected exceedance probability (pxp) across participants, and the overall group-level Watanabe-Akaike weight. Labels are omitted when model indices < 0.1.

